# Seeing or believing in hyperplexed spatial proteomics via antibodies. New and old biases for an image-based technology

**DOI:** 10.1101/2024.08.02.606335

**Authors:** Maddalena M Bolognesi, Lorenzo Dall’Olio, Amy Maerten, Simone Borghesi, Gastone Castellani, Giorgio Cattoretti

**Author notes:** Corresponding Author: Giorgio Cattoretti, MD Pathology, Department of Medicine and Surgery, Universitá di Milano-Bicocca, 20900 Monza (MI), Italy. equally contributed (Maddalena M Bolognesi, Lorenzo Dall’Olio).

## Abstract

Hyperplexed in-situ targeted proteomics via antibody immunodetection (i.e. > 15 markers) is changing how we classify cells and tissues. Differently from other high-dimensional single-cell assays (flow cytometry, single cell RNA sequencing), the human eye is a necessary component in multiple procedural steps: image segmentation, signal thresholding, antibody validation and iconographic rendering. Established methods complement the human image evaluation, but may carry undisclosed biases in such a new context, therefore we re-evaluate all the steps in hyperplexed proteomics. We found that the human eye can discriminate less than 64 out of 256 gray levels and has limitations in discriminating luminance levels in conventional histology images. Furthermore, only images containing visible signals are selected and eye-guided digital thresholding separates signal from noise. BRAQUE, a hyperplexed proteomic tool, can extract, in a marker-agnostic fashion, granular information from markers which have a very low signal-to-noise ratio and therefore are not visualized by traditional visual rendering. By analyzing a public human lymph node dataset, we also found unpredicted staining results by validated antibodies, which highlight the need to upgrade the definition of antibody specificity in hyperplexed immunostaining. Spatially hyperplexed methods upgrade and supplant traditional image-based analysis of tissue immunostaining, beyond the human eye contribution.

**IMPACT STATEMENT:** Staining with multiple biomarkers a single tissue section, multiplex staining, is changing how we examine in-situ normalcy, pathology and the interrelationship of good and bad biological actors. Bioinformatic pipelines developed to deal with high-dimensional datasets such as single cell RNA sequencing or multiparameter flow-cytometry have been adapted to analogous types of data derived from tissue, and co-exist with conventional image analysis tools and human eye-guided image evaluation. We wanted to evaluate if multiplex staining with more than 15 markers (hyperplexed) has comparable sensitivity to conventional image analysis and if these latter analysis tools carry undisclosed biases and should not be used together with hyperplexed staining. We found that the human eye has a reduced discriminative power for grayscale and luminance levels compared to the 8-bit available spectrum, affecting positive signal recognition above the noise. We also found that the granular analytical power of recent bioinformatic pipelines can extract information from images which defy human eye perception and deliver information unattainable with existing image analysis tools for single-stain images.

## INTRODUCTION

In situ antigen detection in tissues via antibody staining, in transmitted (immunohistochemistry; IHC) or fluorescent light (immunofluorescence; IF) is an established tool in science. It is a space structure preserving assay, complementary to other techniques such as in situ transcriptomics [1] or in situ proteomics [2]. It is also complementing all techniques applied to disaggregated specimens, these latter as single cell suspensions (e.g. single cell RNA sequencing [3]; scRNAseq) or homogenates.

In recent years, in situ immunostaining has evolved from a single stain (IHC, the staple tool of diagnostic Pathology) to multiple (from two to seven or more) IF stains, to a much higher number of simultaneous co-stains, typically in excess of a dozen, in what is called high-plex or high-dimensional in situ staining or targeted antibody-mediated proteomics [4].

Recommendations for standardization of the diagnostic use of multiplex stains followed [5], including antibody validation practices.

An analogous progress occurred earlier with flow cytometry (FCM), a technique which employs conjugated antibodies to characterize single cell suspensions [6]. An acceleration of the evolution of the technique was brought by the use of metal-conjugated antibodies and mass spectrometry for detection (Cytometry by time of flight; CYTOF), in lieu of photodetectors and photomultipliers [7]. The evolution of the technique was accompanied by an evolution of the bioinformatic tools required to handle such an increase in data dimensionality to be analyzed [8]. Most of the bioinformatic tools developed for single cell assays (scRNAseq, FCM) have been applied to the analysis of single cells in tissue sections.

Low-plex staining (∼7-10 biomarkers) are increasingly diffuse, partly owing to the popularity of a signal-enhanced technique (Tyramide Signal Amplification or TSA [9]), however, the Image analysis (IA) required for this type of staining does not differ from what is customarily used for single stain images in IHC or IF [10].

What sets apart hyperplexed in-situ targeted proteomics via antibody immunodetection, the method using high-plex (>15) biomarker determination at cellular or sub-cellular resolution in situ, from other low-plex techniques is the use of bioinformatic tools, proper of other single cell assays [11, 12]. Analogously to FCM and scRNAseq, human visual image assessment is minimal or nil for these processes, despite the ground truth data which are tissue images.

IA tools development [13] has accompanied the production of images all along. Interestingly, one of the main concern of scientists using IA is to identify nuclei in sections [14], something Surgical Pathologists do effortlessly every day.

IA has been developed not as a replacement of the human eye but as a companion, particularly for simplified one-protein-at-a-time diagnostic immunostains [15]. However multiplex staining data are intrinsically so complex that deep learning-assisted IA has an increasing role in multiple steps, such as image segmentation [16], data normalization and cell classification [17]. Yet, the microscope’s future evolution in an expert’s view still features eyepieces [18].

We sought to reevaluate the individual components leading to single cell classification via hyperplexed stains, and in particular the role, when present, of a human visual assessment of images in processes such as assay sensitivity, antibody validation, signal thresholding and gating and cell segmentation.

By analyzing a public human lymph node dataset with a custom bioinformatic pipeline, BRAQUE [19], we found that the human eye is dispensable for the analysis of in situ hyperplexed multistainings.

## MATERIALS AND METHODS

### Ethical background

The study has been approved by the Institutional Review Board Comitato Etico Brianza, N. 3204, “High-dimensional single cell classification of pathology (HDSSCP)”, October 2019. Consent was obtained from patients who could be contacted or waived according to article 89 of the EU general data protection regulation 2016/679 (GDPR) and decree N. 515, 12/19/2018 of the Italian Privacy Authority.

### Human specimens

Sentinel lymph nodes (n=5) were extracted from the laboratory information systems of the San Gerardo Hospital by the Authors with clinical privileges and anonymized. Paraffin blocks and sections to be analyzed were selected by a Pathologist after a review of the Hematoxylin and Eosin (H&E) stain. Only archival formalin-fixed, paraffin embedded material (FFPE) was used.

### Histology

Chilled paraffin blocks were sectioned in a rotary microtome (Leica Biosystems, Buccinasco, MI, Italy) at 3 µm, sections were placed in a warm waterbath and collected on charged microscope glass slides. After an overnight oven incubation in an upright position, they were further processed for Hematoxylin & Eosin (H&E), IHC or IF stains.

### Antigen retrieval

Antigen retrieval (AR) was performed placing the dewaxed, rehydrated sections [20] in a 800 ml glass container filled with the retrieval solutions (EDTA pH 8; 1 mM EDTA in 10 mM Tris-buffer pH 8, Merck Life Science S.r.l.,Milano, Italy; cat. T9285), irradiated in a household microwave oven at full speed for 8 min, followed by intermittent electromagnetic radiation to maintain constant boiling for 30 min, and cooling the sections to about 50° C before use.

### Immunohistochemistry

Primary unconjugated antibodies (Abs) were validated for frozen and for FFPE material according to established criteria [21] (see the Supplementary Tables).

For immunohistochemistry (IHC), optimally diluted, validated primary antibodies were applied overnight, washed in 50mM Tris-HCl buffer (pH 7.5) containing 0.01% Tween-20 (Merck) and 100 mM sucrose (TBS-Ts) [22], counterstained with a horseradish peroxidase–conjugated polymer (Vector Laboratories, Burlingame, CA, USA), washed, developed in DAB (Agilent, Santa Clara, CA), lightly counterstained and mounted.

Serial LN sections were immunostained for the AE1-AE3 pre-made cocktail in a Omnis automated immunostainer (Agilent, Santa Clara, CA) with routine same-day protocols.

### Indirect Immunofluorescence

Multiple immunofluorescent (IF) labeling was previously described in detail [20]. Briefly, the sections were incubated overnight with optimally diluted primary antibodies in species or isotype mismatched combinations (e.g. rabbit + mouse, mouse IgG1 + mouse IgG2a, etc.), washed and counterstained with specific distinct fluorochrome-tagged secondary antibodies (Supplementary Tables) [20]. The slides, counterstained with DAPI and mounted, were scanned on an S60 Hamamatsu scanner (Nikon, Campi Bisenzio, FI, Italy) at 20x magnification. The filter setup for seven color acquisition (DAPI, BV480, FITC, TRITC, Cy5, PerCp, autofluorescence/AF) was as published [23]. Additional data are in the Supplementary material.

### Tyramide signal amplification (TSA)

Sections to be processed for TSA were dewaxed, antigen retrieveal was performed as mentioned, endogenous peroxidase was blocked, incubated with the primary Ab overnight and processed as per the manufacturer’s instruction for Alexa Fluor^TM^ 647 Tyramide (cat. N. B40958; Thermo Fisher Scientific, Vedano al Lambro, Italy), a fluorochrome emitting in the red spectrum where tissue autofluorescence is minimal. The Alexa Fluor^TM^ 647 signal was acquired with a 650/13 nm excitation filter, a 694/44 nm emission filter and a dichroic FF655-Di01 filter [24] and could be combined with other fluorochrome combinations except the ones emitting in the 530-570 nm range, where the TSA-Alexa FluorTM 647 product bleeds. All filters are from Semrock, Lake Forest, Ill, USA. Details of the process can be found in the Supplementary Material.

In the double indirect IF-TSA combined staining, TSA was performed first.

### Preparation of immunofluorescent images for single cell analysis

After the stainings were acquired, digital slide images (.ndpi) were imported as uncompressed .tiff with ImageJ (ImageJ, RRID:SCR_003070). Tissue autofluorescence (AF) was subtracted when appropriate as published [20].

### Image analysis

IHC/IF staining quantitation: Fluorescence images were imported in Fiji [25] (RRID:SCR_002285). For area quantification, inverted images were adjusted (Brightness/Contrast command) and thresholded (OTSU). The stained area value was normalized for the total nuclear area value (DAPI). For IHC, the image was color deconvoluted [26] and the DAB image processed as above. Hematoxylin was used for normalization instead of DAPI. Brightness/Contrast, Math transformation (log) and 3D surface plot was used for visualization (see Supplementary methods).

Two public domain IA tools were used for nuclear identification: QuPath (RRID:SCR_018257) [27] and CellPose 2.0 (RRID:SCR_021716) [28]. Details of the setting for IA are reported in the Supplementary Methods.

Adobe Photoshop 2023 (San Jose, CA) (RRID:SCR_014199) and Adobe Illustrator (RRID:SCR_010279) were used for figure layouts.

### Grayscale tone discrimination test of the human eye

Fourteen pathologists, 11 males and 3 females, aged 43 ± 13.8 years (range 29-71), 14.1 ± 13 years into the profession (range 0-43) were asked to log into the Time magazine website https://time.com/4663496/can-you-actually-see-50-different-shades-of-grey/, perform the test and provide the score obtained. Additional information is provided in the Supplementary methods section.

### Bit depth reduction discrimination tests

Twelve pathologists with diagnostic digital pathology experience examined a series of continuous gray shaded bars and full size four-images composites uploaded into NDPserve (Hamamatsu Photonics) via a provided link. The images encompass the various typology of digital images encountered during diagnostic sign-up (Fig S1). The bit depth of each image in the composite was changed from the conventional 24 bit (8 bit times three, 256 colors each) to 6, 5 or 4 bit. via the Adjustments > Posterize command (Adobe Photoshop 2023), then saved in the new format with the original image size and pixel resolution. The 7-bit image was not used for the histology image test, except for the grayscale gradients bars. The percentage of correct scores for each observer, the image type and bit depth was recorded. Additional information is provided in the Supplementary methods section.

### High dimensional analysis with BRAQUE

BRAQUE [19], an acronym for Bayesian Reduction for Amplified Quantization in UMAP Embedding, has been developed for the global analysis of individual cells in tissue sections stained in IF with multiple biomarkers and uses dimensionality reduction algorithms. Is a Python pipeline for automated cluster enhancing, identification, and characterization.

The key procedure of BRAQUE (whose code may be found on Github at https://github.com/LorenzoDallOlio/BRAQUE) consists of a new preprocessing, called lognromal shrinkage. This preprocessing specifically addresses the problem arising from noise due to crossbleed from neighboring cells, in fact, if single cell data are more distinct and discrete on the other hand spatial proteomics markers assume a more continuous distribution with less clear separation among the modalities [29, 30]

In BRAQUE’s preprocessing a mixture of normal distributions is fitted for each of the log-transformed markers, and then each normal component of the mixture is shrunk toward its mean to help further steps counter this continuity and lack of clear separation.

After this crucial step, the markers are standardized and combined in a 2 dimensional latent spaces by the UMAP algorithm. On this embedding space the clustering of cells is performed by HDBSCAN and lastly each cluster is tested for significant markers, which are ranked by effect size to help experts with cell type annotation.

The output consists of multiple clusters, whose numerosity is defined by the size of the smallest cluster (usually not below 0.005% of the cell number or ∼20 cells). Each cluster is defined by A) markers ranked for probability or possibility to identify the cluster, B) a tissue map of the cells belonging to the cluster and C) the expression of a pre-defined set of diagnostic markers for that cluster, compared to the whole population (Fig S1). Each cluster is classified by an expert supervision into cell types.

The HubMap lymph node dataset HBM754.WKLP.262 (doi:10.35079/HBM754.WKLP.262) was downloaded from the HubMap consortium website (https://hubmapconsortium.org/) as a .csv file, thus pre-segmented by the source.

## RESULTS

### The human eye has a biased vision

In image analysis the human eye is required to discriminate signal from noise or background. Published research shows that humans can distinguish about 870 different shades of gray [31], data which are contradicted by Kreit et al. [32], who sets the gray level discrimination in humans at about 30 shades.

Fourteen experienced observers produce a gray discrimination score of 37.8 (SD ±4.77) out of 50 (Fig. 1) which is below the discrimination of 64 gray tones out of 256 (8-bit scale) (see Supplementary Methods) and in keeping with published results [32] and anecdotal annotations in the public domain (see Supplementary Tables).

**Figure 1.**
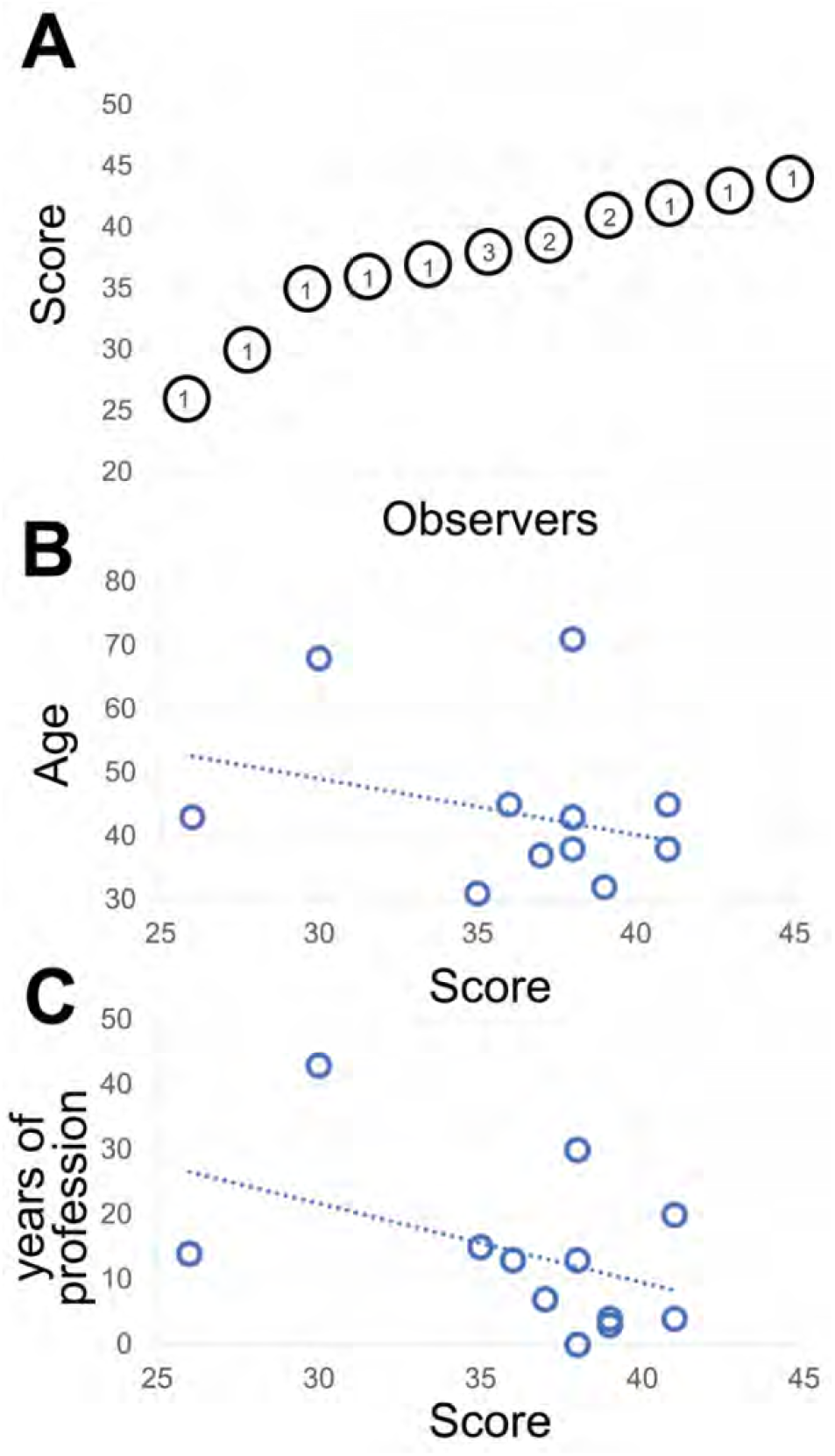
A: The shades of gray score distribution of 15 Pathologists, ordered progressively. The number inside each circle is the number of subjects with that score. B: the relationship between the score (x axis) and the subject’s age (y axis). The R-squared value of the intercept is R^2^= 0.0856. C: the relationship between the score (x axis) and the pathologist’s working experience in years (y axis). The R-squared value of the intercept is R^2^= 0.1876.

The type of images of this test (homogeneously tinted squares surrounded by a thick border) are not the type of images encountered in medicine or biology and may be also prone to hallucinations [33]. We then used microscopy digital images in which the luminance repertoire was reduced from the 256 usual channels (8 bit) down to just 16 (4 bit) (see an example in Fig. 2 and Fig S3).

**Figure 2.**
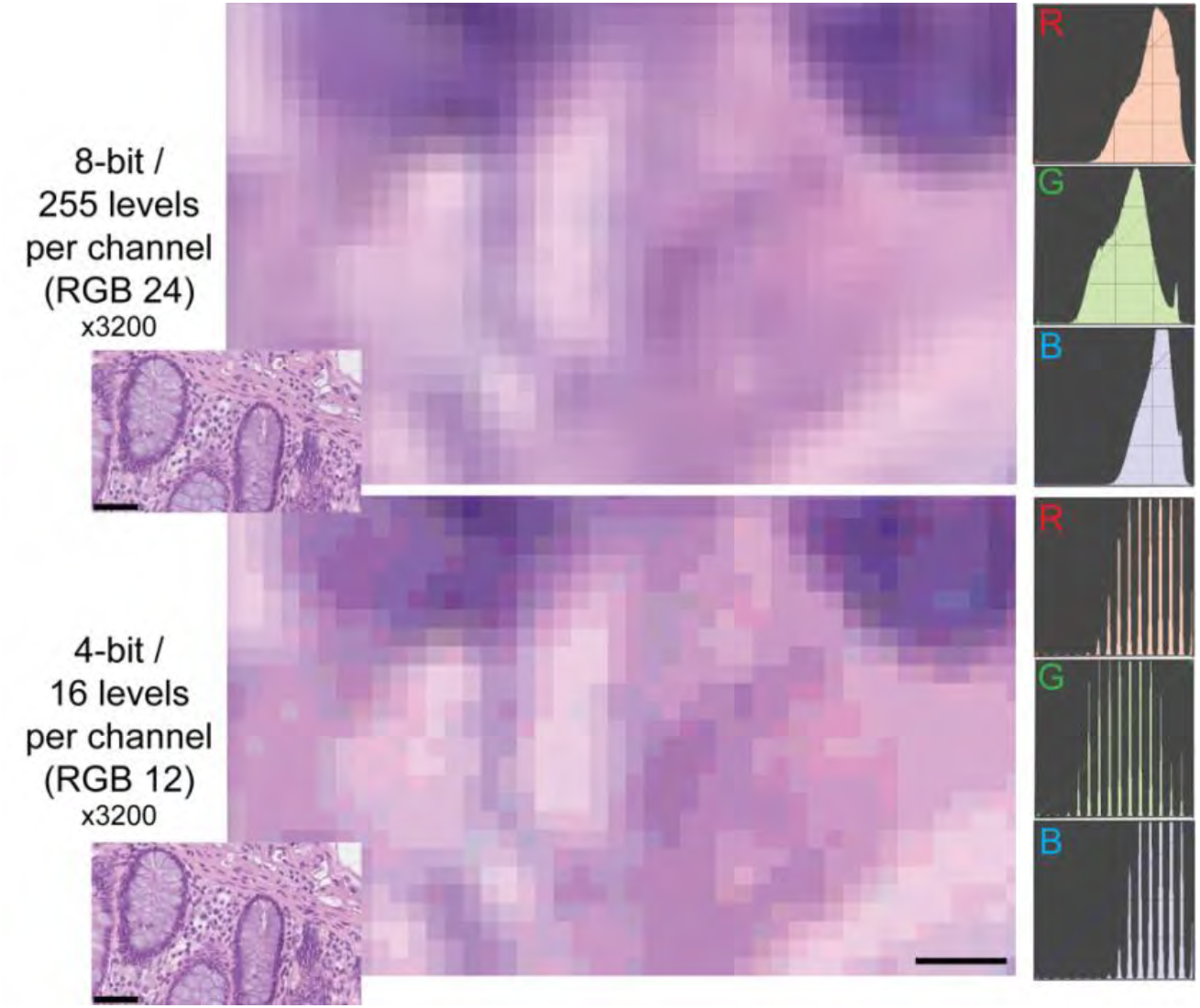
Example of two different bit depths for an H&E image. The two images represent A, a high magnification detail of an 8 bit (24 bits in RGB) H&E image of the human colon, B the same image at 4 bit depth reduction. The R, G and B images on the right are the frequency plots of the image pixels, distributed along the 0-255 channels for each of the three color components. Image A contains 255 levels per channel, image B 16 levels, as shown by the laddering of the RGB profile in the RGB details. Note the same size of the visible pixels.Scale bar: 2.25 µm (five pixels; 3200x). The insets in the lower left corner of each image show the full-size originating images (scale bar 50 µm).

While the observers identify laddering (i.e. reduced bit depth) on the monochrome continuous grayscale images below a mean of 7.7 ± 0.2 bits (range 8-7.5) (Fig. 3A and S3T), they scored correctly the bit depth of only 51% ± 33% of the images (range 26%-75%). There was no apparent relationship between the ability to identify bit degradation in monochrome bands, which scored at the top for all pathologists, and in histology images (Fig. 3A).

**Figure 3.**
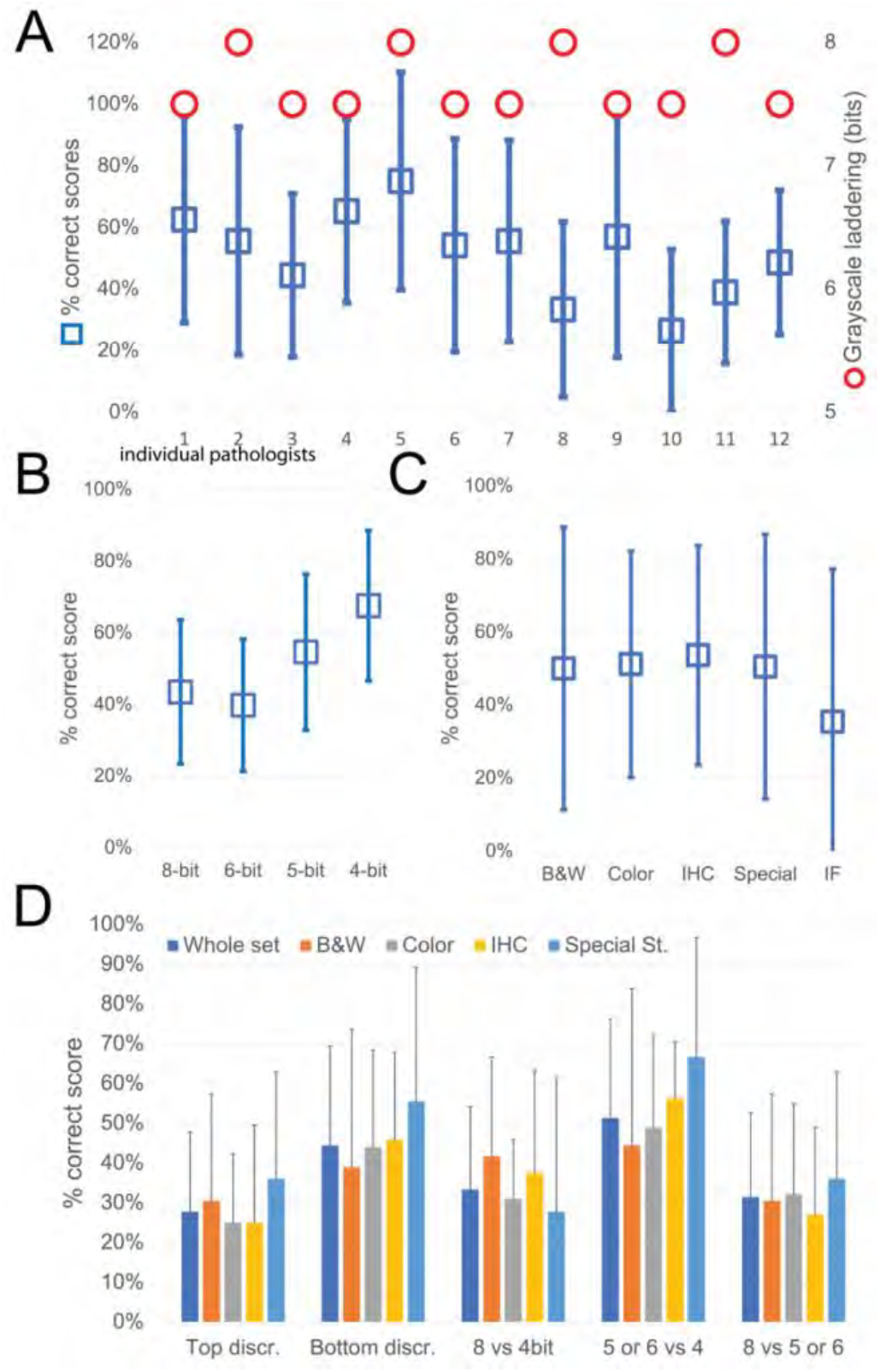
Descriptive graphics of the bit-reduced images scores. A: The two-scale image shows the bit depth below which each of the 12 pathologists identify degradation on a monochrome image (red circles; scale to the right). The mean ±SD percentage of correct bit depth identification on the global test is shown (blue squares; scale to the left). B: Mean percentage ±SD of correct identification on images divided by bit depth. C: Mean percentage ±SD of correct identification on images divided by image type. B&W: grayscale images; Color: H&E-stained images; IHC: immunohistochemistry examples; Special: special stains; IF: immunofluorescence. D: Mean percentage ±SD of correct bit depth identification subdivided between top discrimination (discrimination between 8 and 6 bits), bottom discrimination (5 and 4 bits), 8 bits vs 4 bits, 5 or 6 bits versus 4 bits, 8 bits versus 5 or 6 bits. The scores are further shown for the whole test or divided by image type (B&W: grayscale images; Color: H&E-stained images; IHC: immunohistochemistry examples; Special: special stains)

The most degraded images (4 bit) were more likely to be correctly identified (Fig. 3B). Erroneous bit depth assignment was equivalent in all kinds of common pathology images (Fig. 3C), being a single triple immunofluorescent image the most variably scored (mean 35% ± 42% correct score, range 0%-100%) (Fig. 3).

The discrimination power for degraded images was highest among the bottom range of bit depth (Fig. 3D), with no differences among the image types.

Very detailed images (e.g. colon, testis, LCH) scored marginally better on average than images with low details (brain, muscle, IHC), with 55.8% vs 49.7% correct answers.

From these experiments, we conclude that the discriminative power of the human eye for details along a 8 bit luminance scale is significantly reduced, compared to the available range.

### Signal enhancement methods may deliver marginal gains

Positive signal brightness affects detection. Thus, we wanted to define the sensitivity of the immunofluorescent techniques used in multiplexing, compared to a brightfield standard, DAB IHC. To do so, we used widely used algorithms for immunostain separation from background and identification such as Otsu and K-mean clustering, which are based on vector quantization. These algorithms do not require tuning and the result reflects the image ground truth according to the gray levels of the image.

As previously published by others [34, 35], TSA was not superior, compared to DAB, and as good as double indirect IF [20] for some fluorochromes (Fig. 4). The use of signal-enhancing methods for immunofluorescent staining may marginally benefit signal detection.

**Figure 4.**
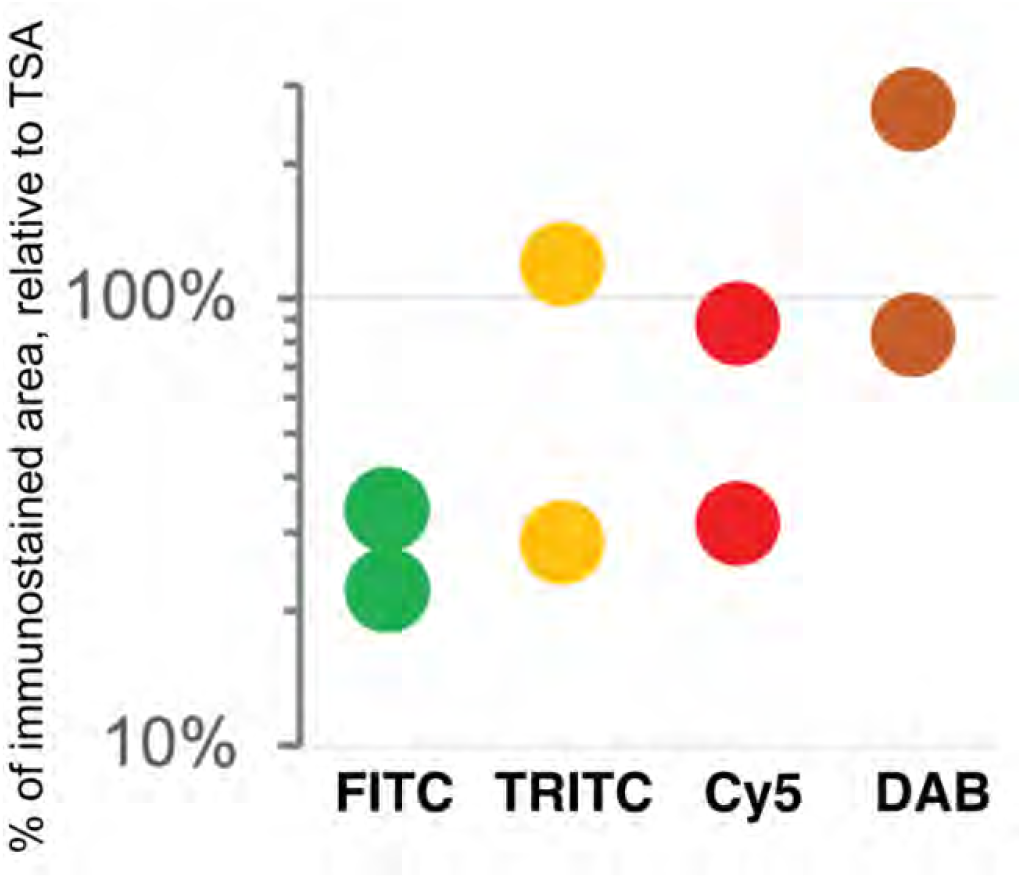
Comparison of sensitivity of detection systems. The area of detection of anti-LMW KRT in serial LN section by secondary Abs conjugated with three fluorochromes (FITC = Alexa 488 green, TRITC = Rhodamine Red^TM^ X orange, Cy5 = Alexa 647 red) and DAB is plotted on a logarithmic scale, relative to the area detected with TSA Alexa 647 (100%). Duplicate experiments.

### Simplified image analysis tools lack sensitivity

For quantification, we used images of abundant low molecular keratin 8 and 18 (LMW-KRT) expressed in thin dendrites of fibroblastic reticular cells (FRC), and we found that commonly used thresholding algorithms to quantify IF immunostains fail to account for positive pixels at the low end of the spectrum, despite being visible to the human eye after image rendering (Fig. 5).

**Figure 5.**
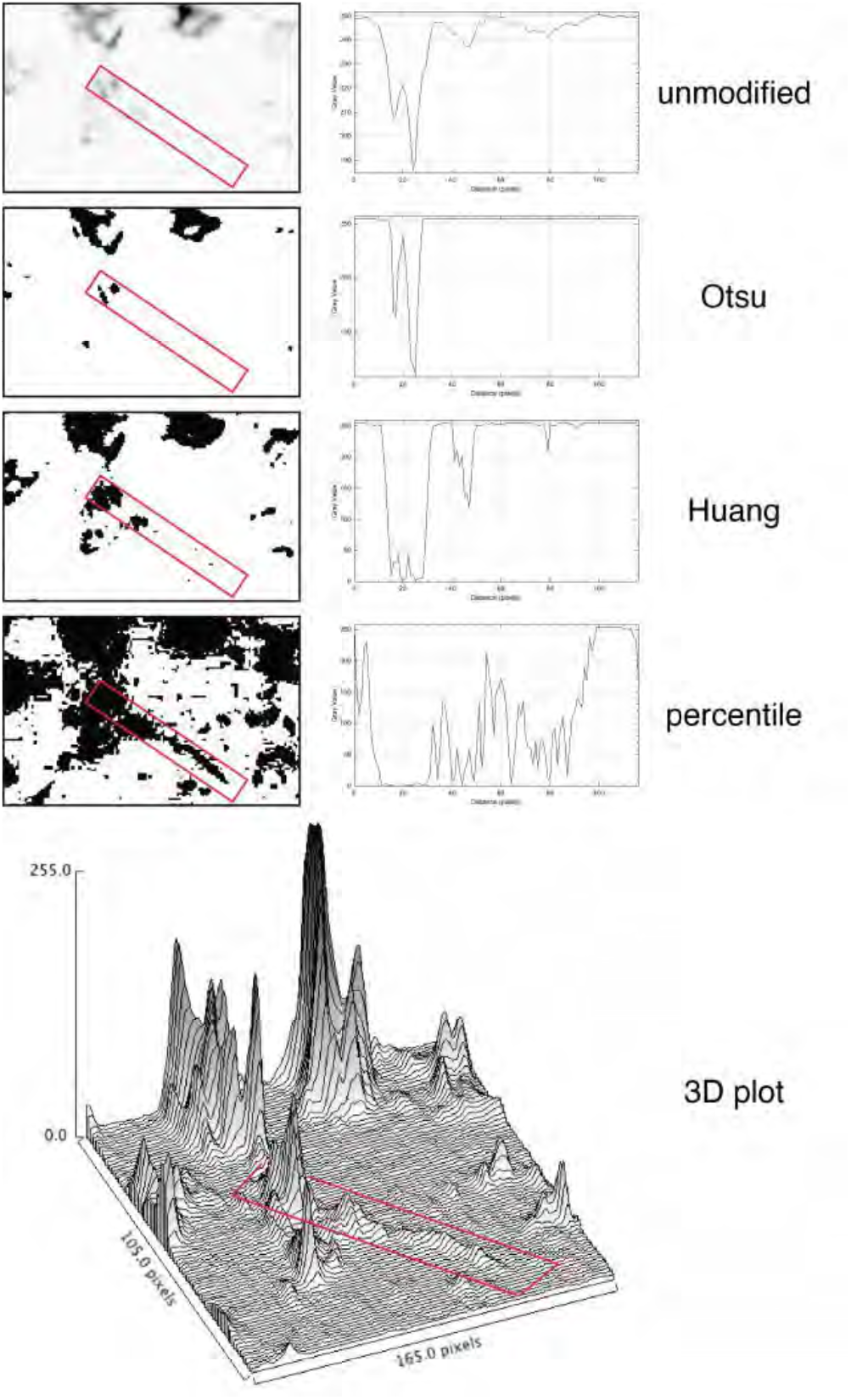
Thresholding LMW KRT staining in LN. The LMW KRT IF stain detail is shown as an inverted, unmodified image, modified with three different Fiji thresholding algorithms (Otsu, Huang and percentile) and as a 3D plot. Only three thresholding algorithms are shown, out of 17 tested. A FRC CK+ dendrite is highlighted with a red rectangle and the mean pixel density value along that rectangle is shown next to the image. Note that, because of the image inversion, daker pixels have lower values in the plot. The continuous intensity variations of the signal above background can be appreciated in the 3D plots. Note that the percentile algorithm highlights numerous background spots in addition to the dendrite of interest. The image shown measures 105 x 165 pixels (47.5 x 74.25 µm).

### Hyperplex staining methods have superior sensitivity

Next we tested the analytical power of hyperplexed stainings by examining a public human lymph node dataset, composed of 28 antibodies (+ DAPI), for which the segmentation method was previously published [36]. In the panel, a widely used “pan keratin” antibody cocktail, AE1 and AE3 [37] was used.

Only AE3 is able to detect KRT8, one of the two LMW-KRT in LN (the other is KRT18). The AE1-AE3 cocktail (“panCK” or “CK”) is used daily by thousands of surgical pathologists to detect nodal metastasis from carcinoma. Because a selective epitope condition prevents the broad detection of LN FRC with this cocktail, only occasional KRT8+ cells are being detected [38]. The presence of a panCK reagent in the dataset made a comparison of the detection power possible between single-stains and hyperplexed images, by enumerating the positive cell types.

We applied to the HubMap dataset BRAQUE [19], a dimensionality reduction algorithms-based analytical pipeline (DRAAP) designed for the spatial Ab-based proteomic data in multiplex, which do not requires pre-definition of positive signal thresholds.

BRAQUE was able to identify 12 clusters containing CK+ cells for a total of 16,698 cells out of 188,450 (9.24%) (Table 1, Supplementary Tables and Supplementary Fig. S1, S2).

**Table 1.** Stromal cells and phenotypes. Phenotype and distribution of stromal cells in the lymph node. Note that all cell types are CD45 negative and devoid of hematolymphoid selective markers. Abbreviations: FRC: fibroblastic reticular cells; SMA: smooth muscle actin; CK: pan-cytokeratin; CollIV: Collagen IV; PDPN: podoplanin; VIM: vimentin; FDC: follicular dendritic cells; For a complete list of cell types, see Supplementary Tables.

| **BRAQUE cell classification** | **clusters** | **total cells** | **%** | **% of total** | **Typical phenotype** |
| --- | --- | --- | --- | --- | --- |
| CK+ FRC | 9 | 13621 | 32.17% | 7.54% | SMA, CK, CD35, CollIV, PDPN, VIM |
| fibroblasts | 5 | 7776 | 18.37% | 4.30% | VIM, CollIV, PDPN |
| myofibroblasts | 3 | 420 | 0.99% | 0.23% | CD34, SMA, CollIV |
| endothelium (CK+) | 3 | 3077 | 7.27% | 1.70% | CD34, CD31, VIM, CK |
| endothelium | 10 | 7021 | 16.58% | 3.89% | CD31, CD34, CollIV, VIM, |
| Lyve1+ endothelium | 8 | 8755 | 20.68% | 4.84% | Lyve1, CD107a, CD31 |
| FDC | 2 | 1671 | 3.95% | 0.92% | CD21, PDPN, CD35, CD20 |
| Total stromal cells | 38 | 42341 | 100.0% | 23.43% |  |
| Total Cells | 90 | 180708 |  | 100.0% |  |

Nine clusters expressed SMA together with CK, a known phenotype of FRC [39], together with variable expression of CollagenIV, CD35 and Podoplanin. These cells accounted for 7.54% of the LN population (13,621 cells).

Three clusters (3,077 cells, 1.70%) had an endothelial phenotype (CD34+ CD31+), where the CK signal could be bleeding from adjacent FRC. 8,196 cells (4.54%) had a stromal phenotype devoid of CK, divided into fibroblasts (4.30%) and SMA-expressing myofibroblasts (0.23%). The tissue distribution of CK+ FRC and fibroblasts is partially overlapping and distinct from endothelial cells (Fig. 6).

**Figure 6.**
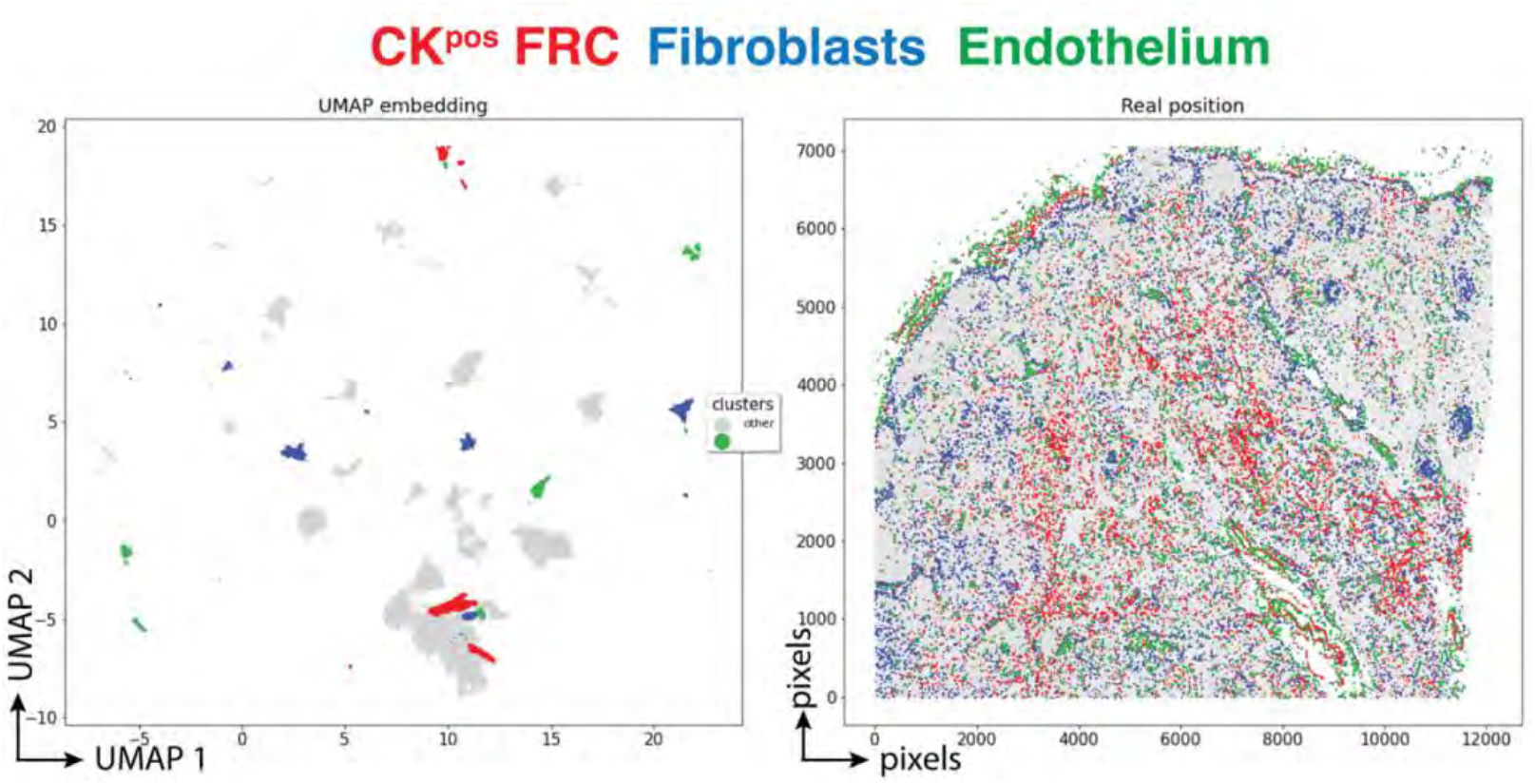
Tissue distribution of FRC and stromal cells. The distribution of CK+ FRC (red; clusters 3, 6, 12, 22, 23, 56, 60, 78, 87), fibroblasts & myofibroblasts (blue; clusters 24, 31, 32, 36, 54, 59, 73, 82) and endothelial cells (green; clusters 13, 14, 17, 25, 28, 29, 37, 46, 53, 72) is shown plotted on the UMAP plot (left) and on a gray image of the LN section (right). The x and y scale on the left are UMAP virtual space arbitrary references, on the right real pixel image dimensions (0.45 µm per pixel). The gray outlines represent the remaining cell clusters (left) and the total of single cells (right).

A distinct population of Lyve1+ sinus lining cells coexpressed CD31, Vimentin, CD107a but not CD34 (Supplementary Fig. S4).

None of these stromal clusters expressed CD44, CD45 or CD45RO, or any other leukocyte restricted marker. The complete cell classification results are available in the Supplementary Methods (Supplementary Tables).

The spatial distribution of the FRC clusters is consistent with the known tissue location and the percentage of total stromal cells (Table. 1).

7.4% of the segmented cells were contained in nine clusters which could not be classified (unclear, artifacts) in addition to cells discarded by BRAQUE (6.8%) upfront on a statistical basis.

By analyzing the same dataset with Phenograph, a similar classification was obtained, including the identification of CK+ stromal cell clusters (Supplementary Fig. S5).

To estimate how the percentage of CK+ FRC detected by BRAQUE in the single HubMap LN would position among the results obtained with an available single-stain, single cell quantitative tool, QuPath, we quantified the IHC stain of two different antibody cocktails: the AE1-AE3 mixture and a two-rabbit monoclonal antibodies cocktail directed at low molecular weight keratin 8-18. AE1-AE3 labelled 9.27% ± 6.79% cells (5 sentinel LN), the LMW keratin cocktail 8.05% ± 7.02% (4 sentinel LN). We also used QuPath to quantify the IF stains used for the TSA and the control experiments. Quantification of IF stains in QuPath was highly erratic because of the difficulty of discriminating by eye signal from autofluorescent background (see Supplementary Tables).

### Hyperplexed staining methods can handle images unreadable by the human eye

We obtained the raw IF images from the HubMap dataset and we could not visualize a distinct CK+ cell population except by applying an image log transformation combined with virtual 3D visualization, after which we were able to identify a weak CK signal co-localizing with vimentin and SM actin (Fig. 7A and Fig. S6).

**Figure 7.**
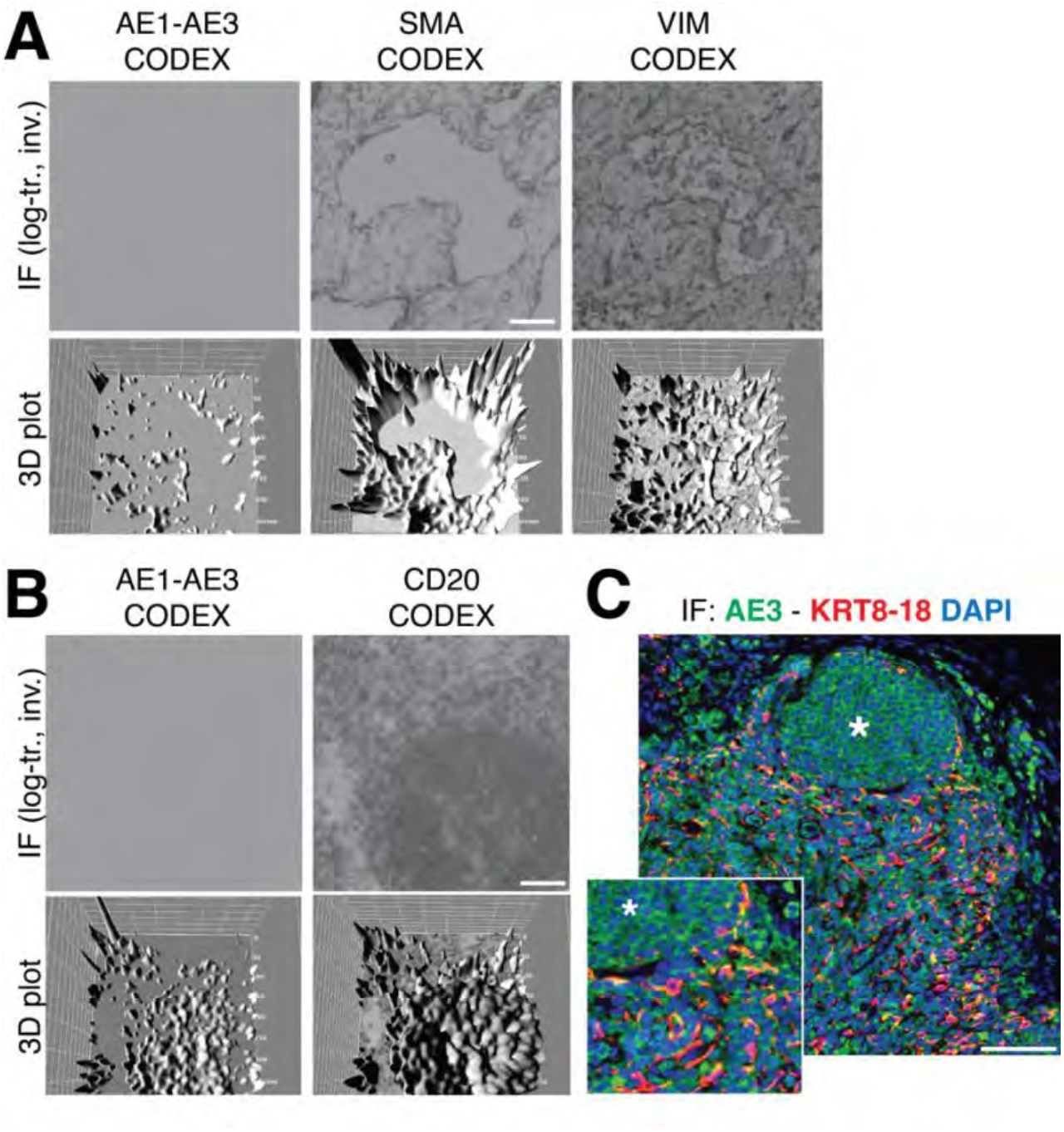
Image rendering of CK+ FRC. A- a detail of three markers, CK AE1-AE3, Vimentin and SMA as 2D log transformed and inverted fluorescence CODEX images (top row) and the 3D transformation of each (bottom row), is shown. B- The same type of images are shown for another tissue detail, comparing CK AE1-AE3 and a B cell marker, CD20. Scale bar: 100µm. C- a low and a high power magnification detail show a three-color image of a LN, produced in-house, where nuclei are blue (DAPI) CK AE3 is green and a LMW KRT cocktail is red. In this image, note a B cell follicle (asterisk) and coexpression of AE3 and LMW KRT in FRC as yellow (green + red). Scale bar: 100µm

Since additional sections from the CODEX LN sample were not available, we used in-house processed FFPE LN sections and stained them with an aliquot of the original AE3 antibody, which we found still effective on a positive control after 37 years [40] (Supplementary Fig. S7). By optimizing the staining conditions [41] and enhancing the image contrast, numerous AE3+ cells co-localize with a LMW KRT staining (Fig. 7C), in addition to other stained cells. The latter appeared to be B cells, based on location and aggregation (no B cell markers were co-stained). TSA amplification of the AE3 signal did not improve the stain (Supplementary Fig. S8, S9).

The staining pattern of in-house CK staining of the lymph node reproduced what was obtained from the CODEX dataset, including the follicular B cell staining (Fig. 7B).

## DISCUSSION

The upper discrimination limit of 64 shades of gray we have shown implies that in the best scenario, signals 4 gray intensity levels (at 8 bit) brighter than noise (256 divided by 64) cannot be discriminated against by the human eye, in addition to known visual and cognitive traps [33]. We thus confirm published data [32] and a number of publicly available anecdotal observations (see Supplementary Tables).

Lack of discrimination of signal from noise broadly affects the appreciation of the full spectrum of biomarker distribution, both in light microscopy and IF. Despite having at disposal the most sensitive stain, IHC with DAB, IA tools are based on algorithms which do not reliably account for signals at the low end of the spectrum. In IHC and IF, a dichotomous image representation (pos/neg) after thresholding is the rule, because of the human eye limit.

Manual gating (i.e. the application of a threshold/barrier discriminating two sets of variables) is considered the main source of data variability and inconsistency in FCM [8, 42, 43], but has never been addressed in tissue staining. The lack of awareness of this limit, results in the vast majority of high-dimensional approach to in-situ cell classification to take advantage of a gating strategy at some point during the process [44–48].

Establishing a signal threshold has three effects, not all negative: *i)* positively assigning a cell to a lineage, *ii)* removing “unwanted” evidence and *iii)* limiting the discovery of new cell types, based on novel phenotypic profiles.

In hyperplexed staining, the limitation in the number of Abs which can fit in a spatial panel forces the selection of biomarkers with *i)* high “diagnostic” value, *ii)* dichotomic expression and *iii)* little overlap with other markers in the panel. A gating strategy to cluster classification [45, 48] is a consequence.

This sparing choice results in a deductive approach to the cell classification, which is not ideal to discover new cell types [44] and prone to overlook unexpected reactivities.

The selection and validation of the antibodies for in-situ staining [49] is made in general either in unrelated substrates (cell extracts, FCM, tumor clonal proliferations) or in tissue staining devoid of phenotypic detail, mostly derived by single color IHC (see Supplemental Data S1 in [48] as an example). Strong staining by visual inspection is favored while concurrent additional weak tissue reactivity is ignored and cataloged as “background”.

Dimensionality reduction algorithms-based analytical pipelines (DRAAPs) extract from spatial images granular data which cannot be acquired by human eye-guided visual representation nor by manual signal thresholding. Both Phenograph and BRAQUE sample the whole range of pixel values, with the difference that BRAQUE provides a phenotypic profile for each cluster, which is based on statistically ranked characterizing markers, chosen from the whole biomarker set, agnostic of the cell type definition meaning and not relying on preset thresholds. The roster of cluster-defining markers includes expected diagnostic Abs, but also unexpected novel expressions such as CD35 and podoplanin (PDPN) on FRC and CD107a/LAMP1 on sinus lining Live1+ endothelial cells.The advantage of BRAQUE and FlowSOM [50] is to provide marker-agnostic cluster-by-cluster evaluation of the key markers statistical relevance (BRAQUE) or mean marker intensity (“star charts”; FlowSOM), and to present the evidence to human judgment.

BRAQUE introduces an innovative data pre-processing step, Lognormal Shrinkage, which is able to enhance input fragmentation by fitting a lognormal mixture model and shrink each component towards its median [19]. It is therefore able to further subdivide signals in the low range and feed these discretized values to the DRAAP. The final effect is somewhat analogous to the introduction of the “logicle” module for FCM [6].

As a result, BRAQUE is ranking AE3 levels in FRC and in those cells only in the significant first or second tiers (see Supplementary Tables) despite the very low levels. In other words, BRAQUE provides statistical strength to the visual perception in Fig. 7C that AE3 and LMW KRT are co-expressed, thus validated according to the “differential antibody” validation criteria [21, 51], but in those cells only. Worth noting that BRAQUE allocates the CK signal in B cells only and not in other hematolymphoid cells (Supplementary Data), and in the lowest tier when significant.

Notably, there is another antibody which unexpectedly shows up in FRC: CD35 (Supplementary Tables and Supplementary Fig. S4). CR1 (the protein name of CD35) is not listed in the Human Protein Atlas (https://www.proteinatlas.org) to be expressed in fibroblasts and in LN, only in the follicular dendritic cells, B cells and macrophages.

CD107a (LAMP1) is listed by the Human Protein Atlas as ubiquitous, however, according to BRAQUE, is differentially expressed only in Lyve1+ endothelial cells and some types of macrophages. These data would not be anticipated by a traditional imaging (Supplementary Tables and Supplementary Fig. S4) or image analysis. Interestingly, BRAQUE do not lists LAMP1 among the ranked markers in neutrophils (Supplementary Data), because the mean expression in these cells falls within the average variation of the rest of the cells.

Our experience from ongoing research (manuscript in preparation) is that the analytical power of DRAAPs and of BRAQUE in particular will discover quite a few other examples of “validated” antibodies which need to be reassessed because of single cell classification. This may be due to inadequate validation upfront, however we favor the hypothesis of missed low-level expression during antibody characterization.

The shortcomings of a visual-guided appreciation of in situ immune detection are numerous. There are published data showing expression of certain biomarkers which have never been reproduced by in situ staining; one example is CD5, shown by RNA and protein on conventional dendritic cells type 2 (cDC2) by high dimensional analysis [52–58] but not on tissue with in situ IF (Wood et al. 1992 [59] and manuscript in preparation). Another example is AID, the enzyme required in the nucleus to perform DNA alterations, which for some time has been detected only in the cytoplasm [60] and still not detected [61] in the presence of the RNA message [62].

Notices about the limitation of an eye-guided approach in high-dimensional studies have just begun to appear in the specialized literature [63, 64].

DRAAPs sensitivity is superior to IA tools used in a conventional setting of low-plex staining. However, saying that DRAAPs are more sensitive is an oversimplification, which blurs the details of how this result is acquired.

First, the algorithm computes the mean expression of all markers in a given cell against all others. Most importantly, data are analyzed as continuous variables, as for FCM [65], because they use normalized mean signal intensity data from single cells, despite the fact that biomarkers may be selected for all-or-nothing expression.

Second, there must be enough biomarkers in the panel in order to classify as different cells which would otherwise be clustered together. Note that DRAAPs can identify cells not only based on present, but also on absent markers [66].

Third, the algorithm must be robust enough not to be disturbed by noise [45].

Fourth, unlike Principal Component Analysis (PCA) which requires at least 2 dimensions, other DRAAPs do not have a minimum number of dimensions to identify meaningful relationships among the data; however, the higher the number of dimensions/parameters provided, the better the discriminative power.

And as a word of caution, fifth, DRAAPs work in a relative space run by mathematics, and can score segmented cells as “negative” for a given biomarker, because statistically below a “mean average” or not above the noise level; in some cases the mean average signal may be considered “positive” by human visual evaluation.

In case the markers are not gated in advance, the product of the DRAAPs is a probabilistic phenotype, because of the inner mathematical working of the algorithm. To go from there to a cell type cluster classification, other steps are required: deep learning cell classification [67] and/or human intervention, neither envisioning visual appreciation of images.

In conclusion, it is about time for hyperplexed spatial proteomics to reduce the dependency from multicolor IF images and the biases associated with human vision and to embrace a space savvy bioinformatic approach like the one that FCM and scRNAseq currently employ. The huge bonus of relinquishing visual imaging and gating is the ability to discover new cell types and cell functions [67], at the cost of revisiting the significance and specificity of the biomarkers which identify such novel populations.

## ACKNOWLEDGEMENTS

The human LN dataset belongs to The Human Body at Cellular Resolution: the NIH Human BioMolecular Atlas Program (doi:10.1038/s41586-019-1629-x). The results here are in whole or part based upon data generated by the NIH Human BioMolecular Atlas Program (HuBMAP): https://hubmapconsortium.org.

The image files and the .csv file for a human LN immunostained with the CODEX platform (now PhenoCycler, Akoya Biosciences, Delaware, USA) are accessed at the HubMap website (https://portal.hubmapconsortium.org/), and for the sample HBM754.WKLP.262 (doi:10.35079/HBM754.WKLP.262) at https://portal.hubmapconsortium.org/browse/dataset/c95d9373d698faf60a66ffdc27499fe1 (accessed last June 11, 2023).

We wish to thank Christopher Wilson (Time Inc. New York, USA) for sharing the code of the website used for testing, Mario R Faretta (IEO, Milan, Italy) for invaluable advice and the Pathologists who contributed suggestions and human visual scoring of synthetic and real-life tests, Elisa Belloni, Francesca M Bosisio, Alessandro Caputo, Giorgio Cazzaniga, Roberta Ciccimarra, Vincenzo L’Imperio, Fabio Pagni, Davide Seminati, Claudio Tripodo, Matteo Zoboli.

## FUNDING STATEMENT

GCat and MMB received funding from Regione Lombardia POR FESR 2014–2020, Call HUB Ricerca ed Innovazione: ImmunHUB. GCast received funding from the EU Horizon 2020 programme (GenoMed4All project #101017549, HARMONY and HARMONY-PLUS project #116026), and the AIRC Foundation (Associazione Italiana per la Ricerca contro il Cancro; Milan, Italy; projects #26216.

## AUTHOR CONTRIBUTIONS

LDA, SB and GCast generated BRAQUE, GCat and AM performed immunostains, MMB, GCat and LDA analyzed the data, GCat and GCas wrote the manuscript.

## LIST OF ABBREVIATIONS

Ab: Antibody
DAB: diaminobenzidine
BRAQUE: Bayesian Reduction for Amplified Quantization in UMAP Embedding
MILAN: Multiple Iterative Labeling by Antibody Neodeposition
scRNAseq: single cell RNA sequencing
IF: immunofluorescence;
FCM: flow cytometry
CYTOF: Cytometry by time of flight
IHC: immunohistochemistry
CK: cytokeratin
LMW-KRT: low molecular weight keratins
FRC: fibroblastic reticular cells
DRAAP: dimensionality reduction algorithms-based analytical pipeline
TSA: tyramide signal amplification
FFPE: formalin-fixed, paraffin embedded
FDC: follicular dendritic cells

## COMPETING INTERESTS

Competing Interests: all the Authors declare none.

## LIST OF SUPPLEMENTARY DATA AND DATA AVAILABILITY

Supplementary Data file (pdf): contains Supplementary Materials and Methods, Supplementary online data availability, Supplementary Figure legends.

Supplementary Tables are provided an Excel file (SupplementaryTables.xlsx).

Additional data with the clusters definition produced with BRAQUE and .ndpi IF images are deposited in Bicocca Open Archive Research Data (BOARD).

Bolognesi, Maddalena; Dall’Olio, Lorenzo; Maerten, Amy; Borghesi, Simone; Castellani, Gastone; Cattoretti, Giorgio (2023), “Seeing or believing in hyperplexed spatial proteomics via antibodies.”, Bicocca Open Archive Research Data, V1, doi: 10.17632/kmxz7fgydx https://data.mendeley.com/datasets/kmxz7fgydx/1

## SUPPLEMENTARY DATA

### Gray tones discrimination test

The website https://time.com/4663496/can-you-actually-see-50-different-shades-of-grey/ provides an evaluation of the discriminative power of the human eye for shades of gray.

A selection of eight new images are presented each accession, containing nine randomly assorted squares, over a background tone. Overall, gray tones ranging from 32 to 228 pixels are shown at any accession, the background tone ranges from 44 to 220 pixels, each 9-square screen displays gray levels, distant 47.9 ±15.2 pixels from the background (88-16), in 4 pixels increments/decrements from the background. These data derive from the analysis of five different tests.

A perfect score (all squares different from background selected and none of the ones identical to the background) is 50. Given these details, a score of 50 (obtained by none of the participants) requires the human eye to discriminate 2-5 background-toned squares among nine for 8 times in a row, whose gray tones are 4 pixels apart. This equals to a discrimination of 256/4 = 64 tones, which may be the top discriminative power of the human eye.

The technical characteristics of the computer screens used were not standardized.

### Bit depth reduction discrimination test

Three micron sections were obtained from anonymous formalin-fixed, paraffin embedded surgical specimens, placed on glass slides and routinely processed for Hematoxylin and Eosin or for immunohistochemical markers in dedicated autostainers.

The Hamamatsu NanoZoomer S60 scanner (RRID:SCR_023762) (Hamamatsu Photonics K.K., Hamamatsu City, Japan) is equipped with a Nikon 20× or 40x/0.75 PlanSApo objective and a linear ORCA-Flash 4.0 digital CMOS camera (Hamamatsu Photonics K.K.).

Slides were scanned at 20x or 40x at a resolution of 0.45 and 0.25 µm/pixels respectively. Selected cases were acquired as 5-stacks zeta-stack and merged in a single-layer image via the Hamamatsu toolkit.

The whole slide images (WSI) were saved as .ndpi files at a resolution of 300 dpi.

Selected areas were opened with the NDP View 2.0 software (Hamamatsu Photonics) at the optical maximal magnification and saved at the original resolution (300 dpi) as .jpg images.

Still life images (as controls) were obtained with a Panasonic DMC-TZ100 camera equipped with a 20.1 megapixel MOS sensor. Images were saved as 16 bit RW2 files.

A bar with a continuous grayscale gradient was created in Microsoft Powerpoint (RRID:SCR_023631) and exported as a 2550 x 556 pixel image, 225 dpi.

Images were imported in Adobe Photoshop 25.0.0 (RRID:SCR_014199) as 8 bit per channel (RGB; 24 bits) and downsized to 7, 6, 5 and 4 bits via the Adjustments > Posterize command, then saved in the new format with the original size and resolution. The 7-bit image was not used for the test, except for the grayscale gradients bars.

Note that the pixel resolution did not change during this process.

An example of the bit depth degradation (or posterization) is shown in Fig. 2.

Bit channel depth will be defined throughout this manuscript referred to a single channel, i.e. 8 bit for maximal resolution (equals 24 bits in RGB channels).

Composite images and labeling were obtained with Adobe Illustrator (RRID:SCR_010279).

Full size four-images composites for scoring were uploaded into NDPserve (Hamamatsu Photonics) and a link provided for image visual analysis. The NDPserve image can be zoomed 2x on screen.

A low-power overview of the image panels is shown in Fig. S3.

Each observer was requested to assign the correct bit depth to each of the four images in each panel, via a pull-down fixed menu. One H&E and one IHC image was scored twice, adding a request to identify similar images among the 4-image panel.

For the continuous gray shaded bars, the observer would identify the bars showing laddering. All pathologists scored all the images.

A fillable PDF scoring sheet (Adobe Acrobat Pro RRID:SCR_023361) with links to the full size images and pull-down scoring menus was provided to each participant. The PDF sheet contains a scoring result summary pages which can be exported to a spreadsheet for analysis.

The type of monitor used and resolution was recorded (see Supplementary material), but not used in the analysis.

The original images and the PDF test are available at Bicocca Open Archive for Research.

All the tests were performed on fully anonymous human specimens, in compliance with the Helsinki declaration for international guidelines for research. Steenbras (Lithognathus mormyrus) was purchased for human consumption at a local market, thus exempt from IRB. Flowers were a gift from a close relative. Non-pathology images were provided as familiar visual references for posterization.

### Public CODEX data

The image files and the .csv file for a human LN immunostained with the CODEX platform (now PhenoCycler, Akoya Biosciences, Delaware, USA) are accessed at the HubMap website (https://portal.hubmapconsortium.org/), and for the sample HBM754.WKLP.262 (doi:10.35079/HBM754.WKLP.262) at https://portal.hubmapconsortium.org/browse/dataset/c95d9373d698faf60a66ffdc27499fe1 , selecting, in sequence: drv_CX_20-008_lymphnode_n10_reg001 > processed_2020-12-2320- 008LNn10r001 > the folder “stitched” > the folder “reg001” and then downloading the images as 16-bit .tiff (accessed last June 11, 2023).

### Tyramide amplification (TSA)

Semi-serial sections from sentinel lymph nodes were selected for sensitivity comparison experiments.

Dewaxed, antigen retrieved FFPE sections were incubated for 10 min. with a H_2_O_2_ - NaN_3_ mixture [68] in order to block endogenous peroxidases. Afterward, the sections were washed in TBS-Ts and incubated overnight at RT with two primary antibodies, mismatched for species.

The next day, the sections were washed twice for 15 min. In TBS-Ts without NaN_3_.

A blocking horse serum (Vector, Burlingame, CA) was applied for 10 min., rinsed once, and a secondary antibody HRP-conjugated polymer (Vector) was applied for 30 min.

After two 15 min. rinses in azide-free TBS-Ts, a freshly made TSA solution was applied for 20 min.. The TSA solution was made of 1 ml of Tris buffer pH 7.4, 10 µl of 100x H2O2 solution (made of 50 µl H2O2 3% in 1 ml distilled water) and 10 µl of the 100x Tyramide stock solution. The Tyramide stock solution was made by dissolving the lyophilized product in DMSO as specified by the vendor (see Pub. No. MAN0015834, Invitrogen) and kept at -20 C°.

The TSA reaction was stopped by several rinses in TBS-Ts with NaN_3_, then the second primary antibody was counterstained for 30 min. with an appropriate secondary antibody. The staining cycle was repeated twice [20]. After a final 30 min. wash, the section was counterstained with DAPI, mounted and scanned.

### Fluorescence acquisition

The Hamamatsu NanoZoomer S60 scanner (Hamamatsu Photonics K.K., Hamamatsu City, Japan) is equipped with a Nikon 20×/0.75 PlanSApo objective, a Fluorescence Imaging Module equipped with a L11600 mercury lamp (Hamamatsu Photonics K.K.), a linear ORCA-Flash 4.0 digital CMOS camera (Hamamatsu Photonics K.K.) and two six-position filter wheels, one for excitation, the other for emission filters, and a three-cube turret. The filter setup is composed of a set of excitation filters, housed in one wheel: 387/11 (DAPI), 438/24 (BV480), 480/17 (FITC), 556/20 (TRITC), and 650/13 (Cy5); one set of emission filters: 435/40 (DAPI), 483/82 (BV480), 520/28 (FITC), 617/73 (TRITC), and 694/44 (Cy5). Dichroic filters are housed in three OM cubes: a triband FF403/497/574-Di01 (DAPI/FITC/TRITC) and two single pass FF458-Di02 (BV480) and FF655-Di01 (Cy5). All filters are from Semrock, Lake Forest, Ill, USA.

AlexaFluor 657 Tyramide was acquired for 96 milliseconds. Double indirect IF acquisition time ranged from 192 to 320 ms for Cy5 and 112 to 192 ms for FITC.

### Image manipulation for viewing

Raw .tiff uncompressed files were produced by a Fiji plugin (Bio-format importer) from .ndpi grayscale fluorescence images.

Two types of image manipulations were used: log transformation and Brightness & Contrast. For log transformation, the math>log command was applied to the dark background IF images, which were then inverted.

The Brightness & Contrast command was set to “auto” to inverted IF images.

Although both methods produce equivalent results, we found the log transformation the best one for images where the positive signal is most minuscule. For composite color images, both the log transformation and the Brightness & Contrast commands were used and adjusted visually.

The “Analyze” command “3D surface plot” was applied to modified IF images and the “Min” and “Max” cursors adjusted manually.

### IHC/IF quantification

Image thresholding was used for IHC or IF quantification. The default setting for the thresholding algorithms in ImageJ/Fiji (https://imagej.net/plugins/auto-threshold) was used. IHC immunostains of cytokeratin-positive FRC and of IF stains were quantified as single cells with QuPath (RRID:SCR_018257); Hematoxylin OD density was adjusted between 0.1 and 0.01, the nuclear outline was expanded by 0.1 µm, cytoplasmic DAB mean OD density was set at 0.1. Additional information can be found in the Supplementary Tables.

### BRAQUE pipeline

BRAQUE is a python pipeline for automated cluster enhancing, identification, and characterization. Its code and documentation (comprehensive of usage instructions) can be found at https://github.com/LorenzoDallOlio/BRAQUE. Such pipeline can either be run on a tidy dataset in .csv format or its functionalities can be cherry-picked to be inserted into different pipelines. The HubMap dataset .csv was loaded and run after minor adjustments.

### Phenograph pipeline

The HubMap dataset .csv was analyzed with a Phenograph pipeline as published [69]. Intensity data for each biomarker and for each cell in the cluster are shown as hierarchical clustering plot (see [69]); the custom R script can be downloaded at https://board.unimib.it/datasets/46r9hcvpbd/1. Note that mean intensity rendering as an heatmap may be deceiving [70].

## SUPPLEMENTARY DATA AVAILABILITY

Data are deposited in Bicocca Open Archive Research Data (BOARD)

## SUPPLEMENTARY TABLES LEGENDS

One single Excel file contains the following tabs:

Primary and secondary Abs: Primary and secondary antibody specifications

Table-allCellsClassified: Classification of all BRAQUE clusters in the HubMap lymph node Table-stromalCells: Classification of all BRAQUE stromal cell clusters in the HubMap lymph node

QuPathMeasurements-IHC_IF: Measurements and settings for QuPath 50shades-Survey: Observer’s scores for the "50 shades of gray" website

PathologistsScoring: Scoring sheets and biometric info for all pathologists scoring bit-degraded images

BitDepth-chahnnelsEquivalence: Chart with equivalence calculations between bits, channels and colors.

VisionLimitsReferences: References to limitation of human visions, else than published papers.

---

**Fig. S1.**
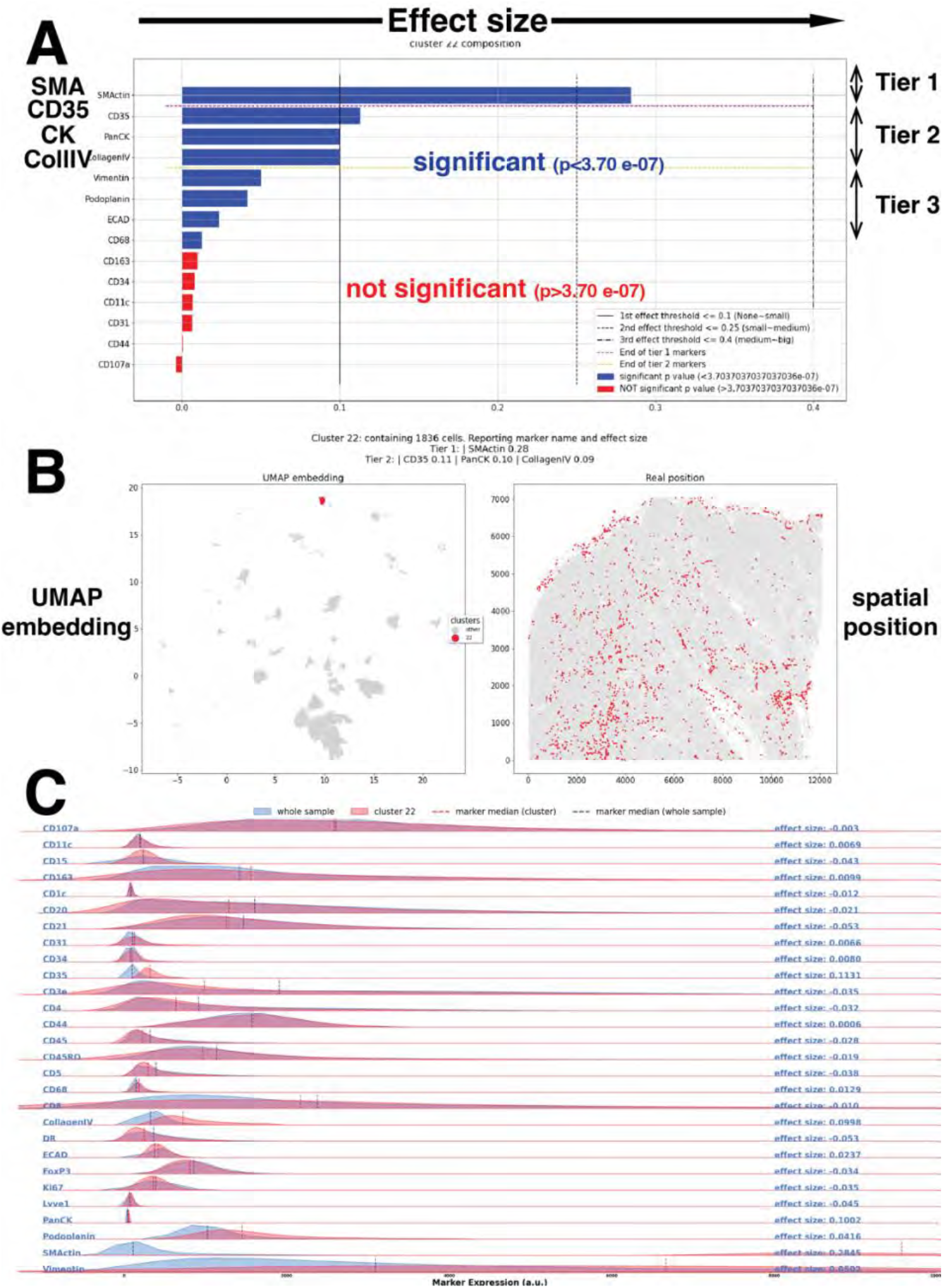
BRAQUE display of an example of a FRC cluster (cluster #22). **A** The markers defining cluster 22 are ranked according to the robust effect size metric *d_signed_*, and colored according to their Welch t-test p-value (blue: significant; red: non-significant). Note pan CK positioned in the Tier 2 position (“possibly” defining). **B** the position of Cluster 22 in the UMAP graph is shown on the left; the cells composing this cluster are superimposed as red dots on an image of the lymph node in grey, on the right. **C** the distribution of 26 markers in cluster 22 (red) is compared with the distribution on the whole LN (blue) in real values.

**Fig. S2.**
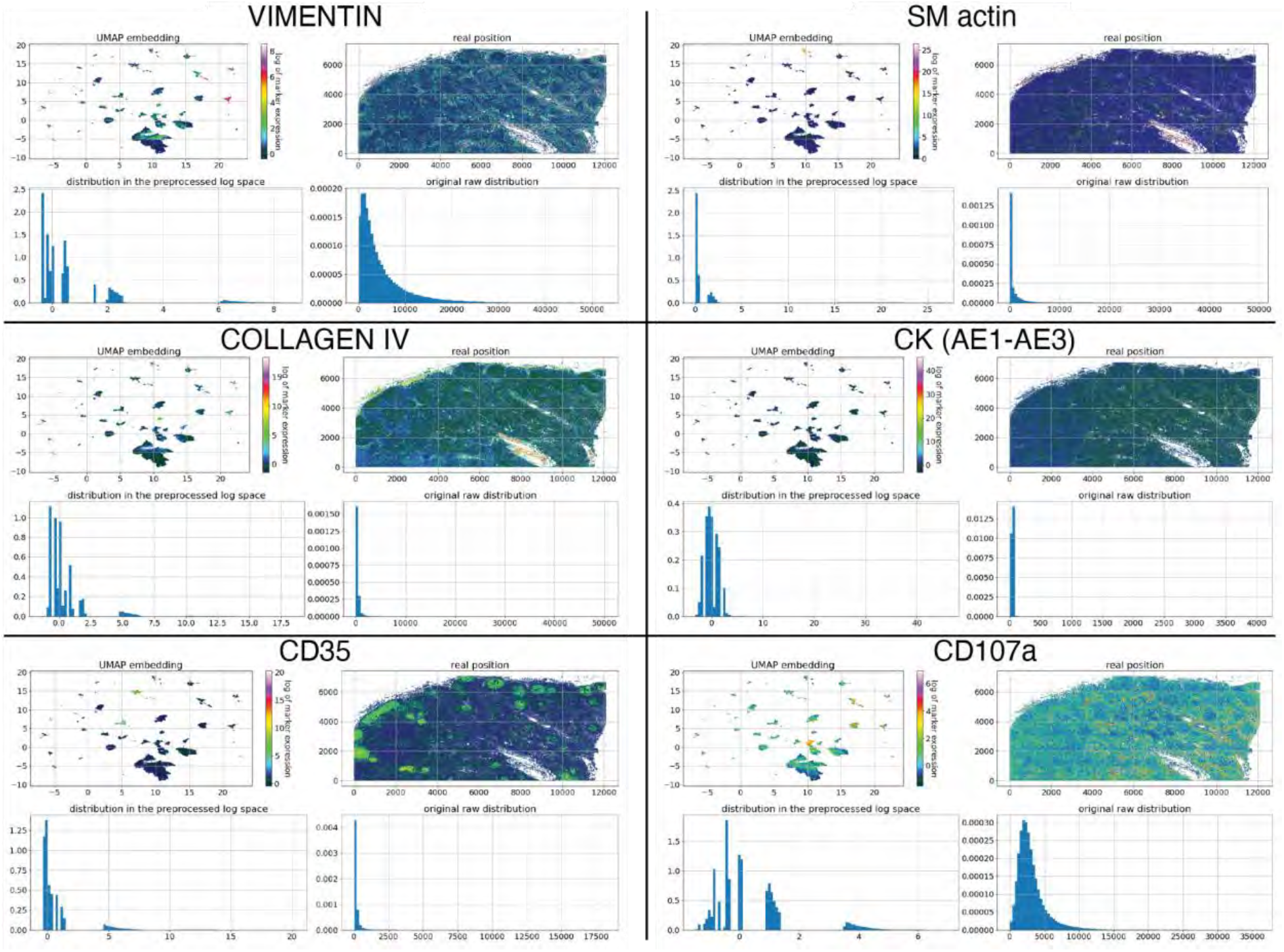
Expression plots and distribution for FRC markers. BRAQUE display of six markers relevant for FRC classification. For each marker, in clockwise order, is presented: the marker log expression according to a color palette in the UMAP plot; the pseudocolor tissue distribution; the original raw intensity distribution; the distribution in the preprocessed log space after gaussian mix distribution and shrinking. Note the continuous data nature in the original raw distribution and the discretization in the preprocessed log space.

**Fig. S3.**
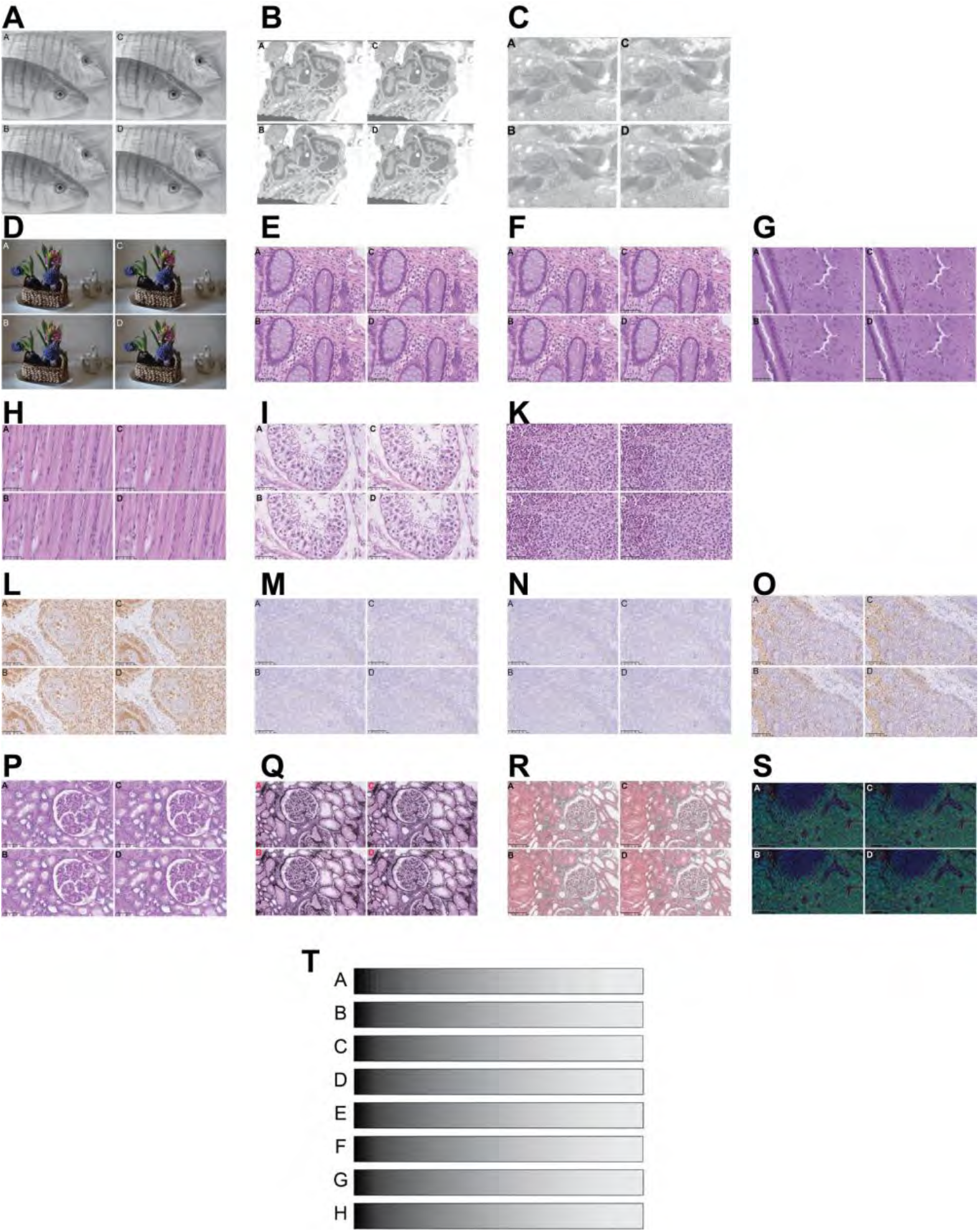
Overview of the image panels tested. Each subpanel consists of four images of different bit depth, the order scrambled for each set, to be scored online full-size. A: steenbras; B: electron microscopy image, low power; C:electron microscopy image, high power; D: flower pot, hyacinths; E, F: normal human colon; G: brain; H: muscle; I: testis; K: Langerhans cell histiocytosis (LCH); L: CD3 IHC on tonsil; M, N: CD30 IHC; O: CD68; P: PAS stain, kidney; Q: Jones stain, kidney; R: trichrome stain, kidney; S: triple immunofluorescence for CD4 (FITC), FOXP3 (TRITC) and DAPI, tonsil; T: monochrome bands, degraded with controls, scrambled.

**Fig. S4.**
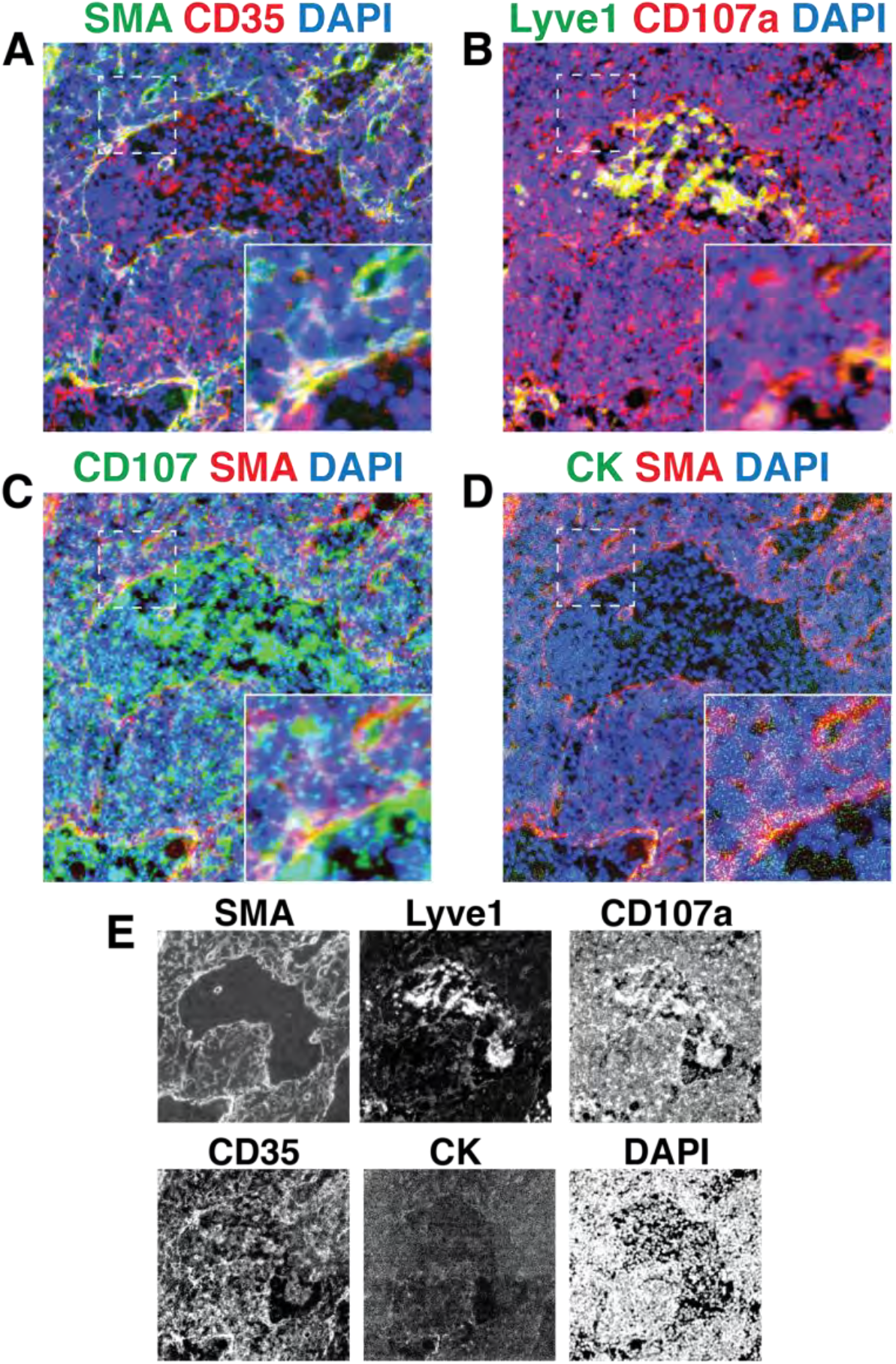
Composite images for coexpression of CD35 or CD107a on FRC and Lyve1 endothelial cells. A detail of the .tiff images for SMA, Lyve1, CD107a, CD35, CK and DAPI is superimposed in color composite. A detail highlighted by the dotted square is in the lower right corner of each image. **A**: SMA and CD35 are coexpressed (yellow) in FRC. **B**: Lyve1 and CD107a are coexpressed (yellow) in endothelial cells and in sinus macrophages. **C**; CD107a and SMA are juxtaposed, but do not colocalize, retaining the green or red color. **D**: CK staining is diffuse and dust-like over the field, occasionally co-localizing with SMA+ cells. **E**: greyscale single stain images of the composite components. All images have been adjusted with Fiji for brightness and contrast in order to be visible to the human eye. Scale bar: 100 µm

**Fig. S5.**
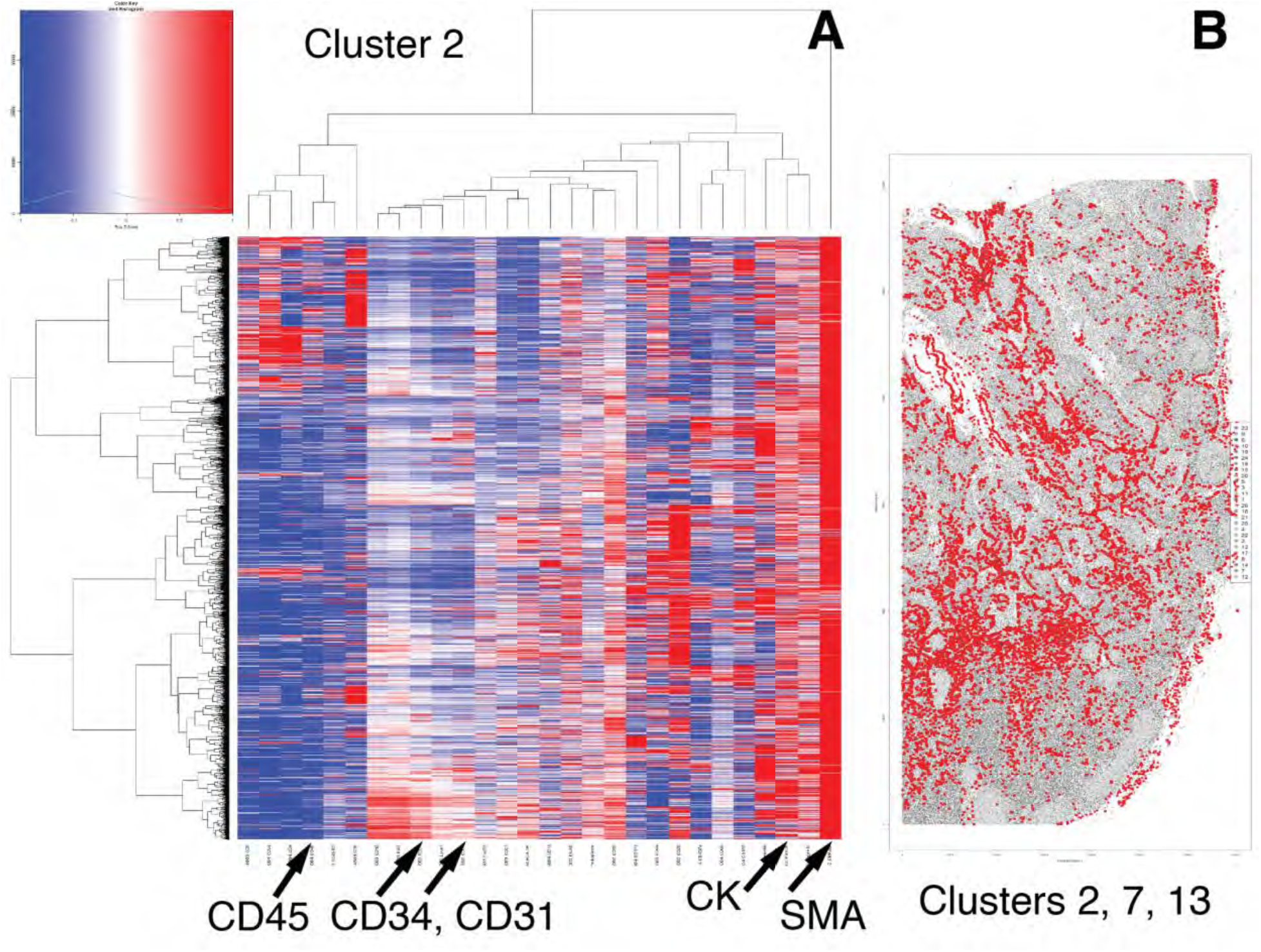
Representative CK+ FRC cluster and tissue distribution by Phenograph. **A**- a representative CK+ Phenograph cluster (cluster 2) contains SMA+ and CK+ cells, which are characterized by negative CD31, CD34 and CD45 expression. Rows are cells, columns are markers, ordered by hierarchical clustering. **B**- CK+ FRC clusters (2, 7 and 13) are plotted in red onto a gray image of the LN.

**Fig. S6.**
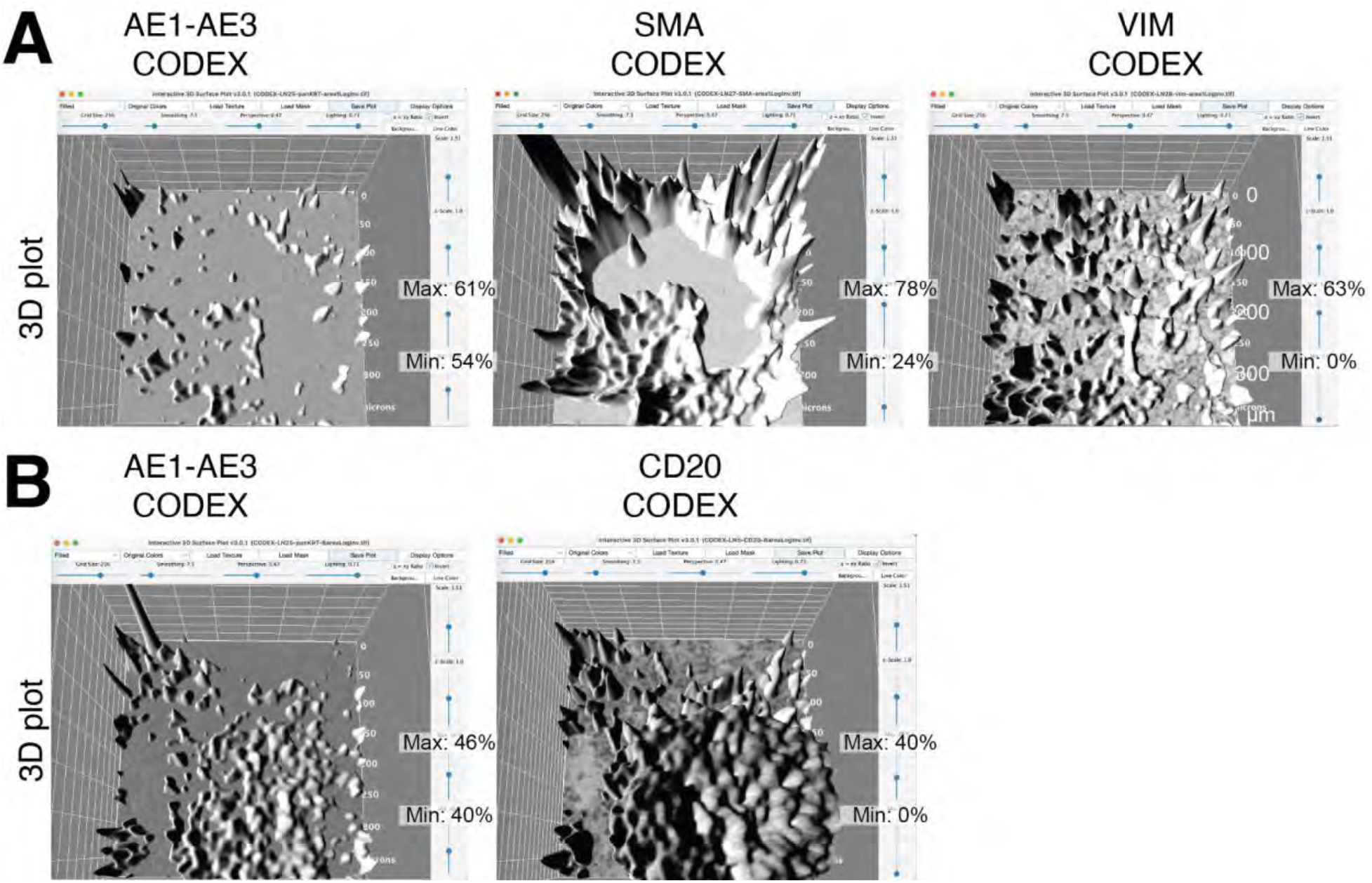
The settings for the 3D plots of Figure 6 are listed for each image. Only setting variating in each image are magnified. Invariant settings on the x axis are: Grid Size: 256; Smoothing: 7.5; Perspective: 0.47; Lightning: 0.71; On the y axis: Scale: 1.51; z-scale: 1.0. The grid white numbers are microns. Note the individual manual adjustments in Min. and Max. values for each biomarker.

**Fig S7.**
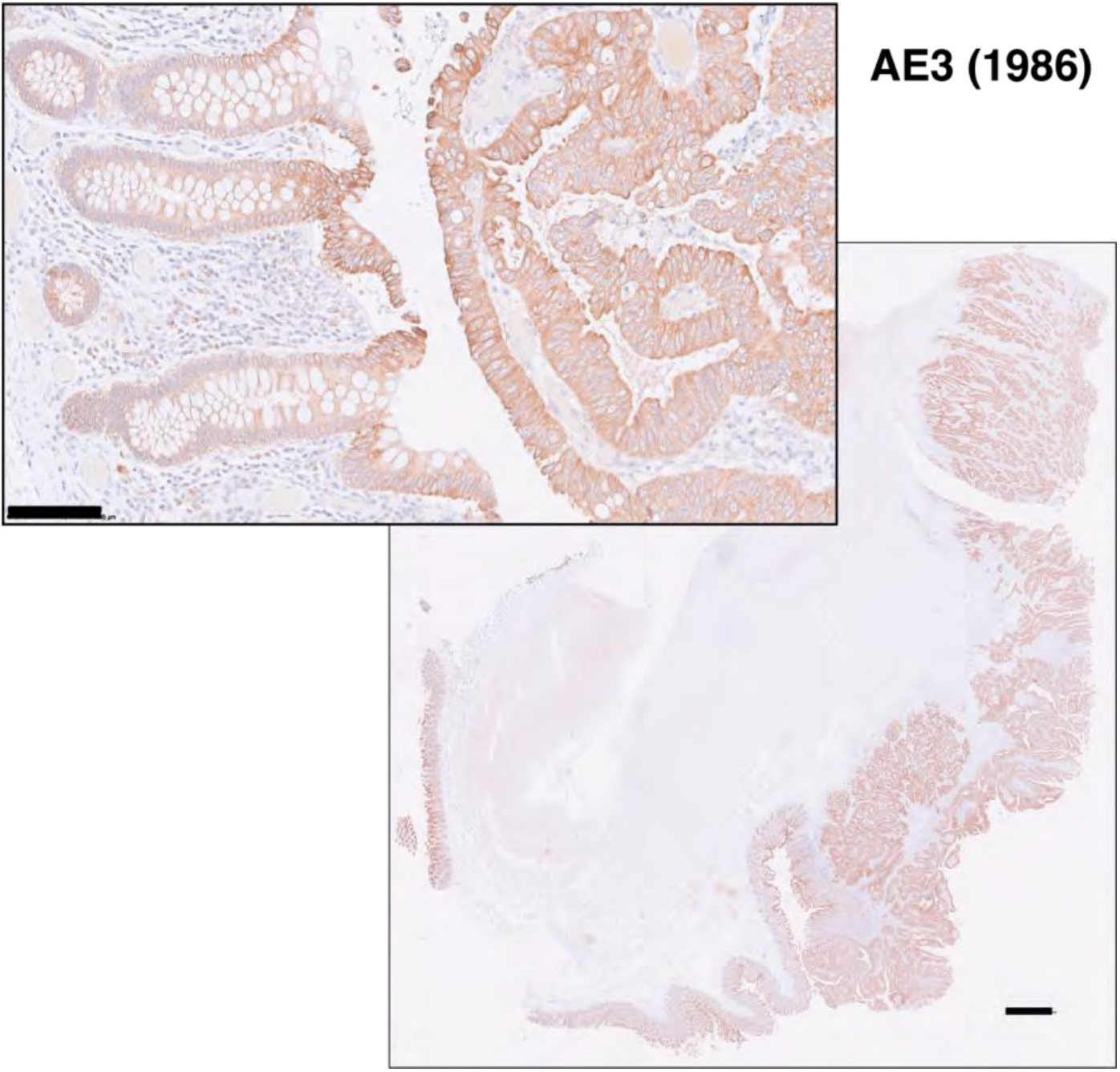
Indirect immunoperoxidase staining (DAB) of a colon cancer specimen with the AE3 antibody obtained in 1986. DAB color is brown, hematoxylin nuclear blue counterstain. Scale bar for the low-power image: 1 mm; for the high magnification inset: 100 µm.

**Fig. S8.**
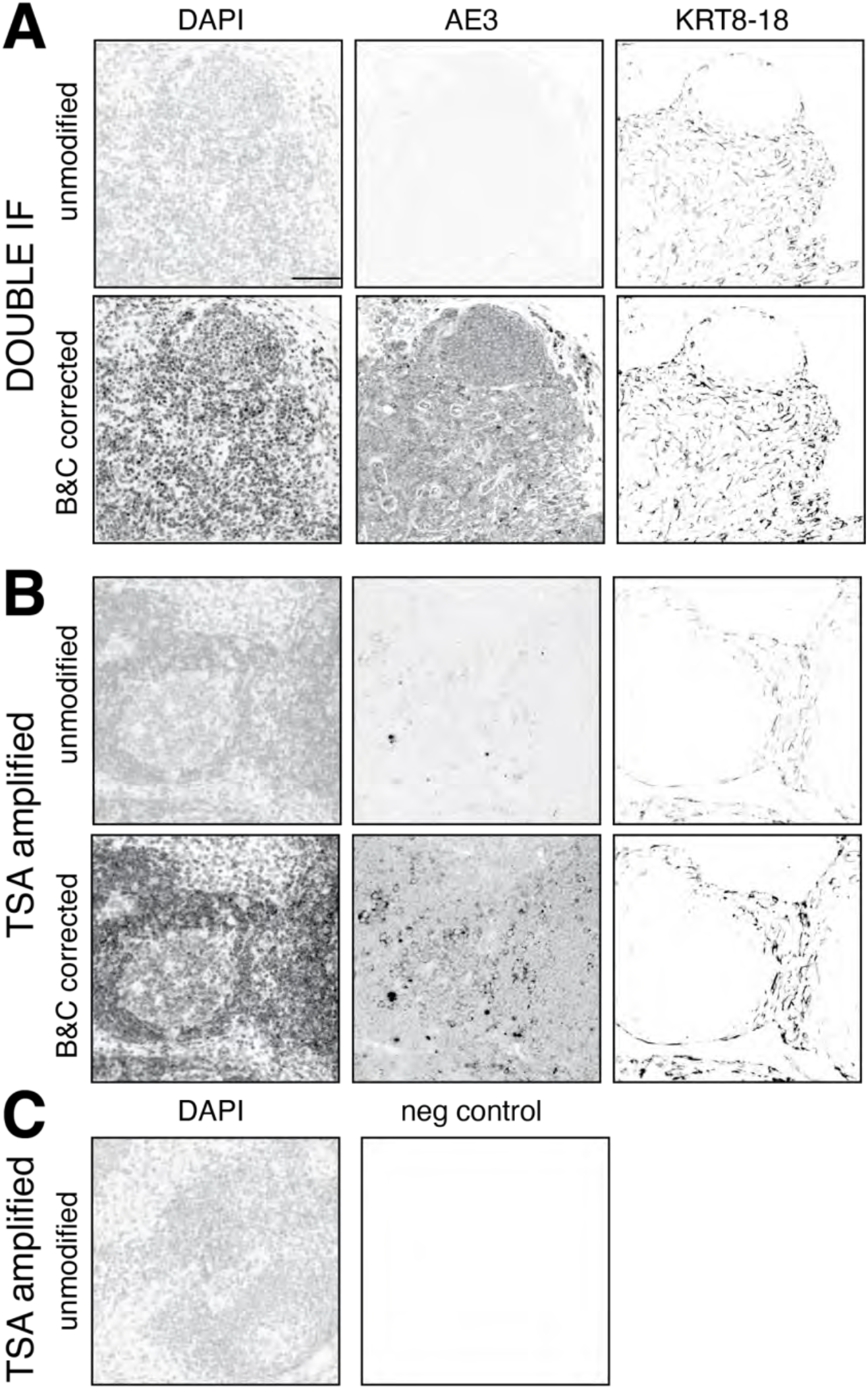
Comparison of staining efficiency and image manipulation for CK staining. CK AE3 and LMW KRT pool staining by double indirect IF (**A**) and by TSA (**B**) are compared. For each method the unmodified (top) and the Brightness and Contrast modifications of inverted IF images (bottom) are shown. **C** A negative control after TSA is shown, unmanipulated. For the image manipulation parameters, see Fig. S9.

**Fig. S9.**
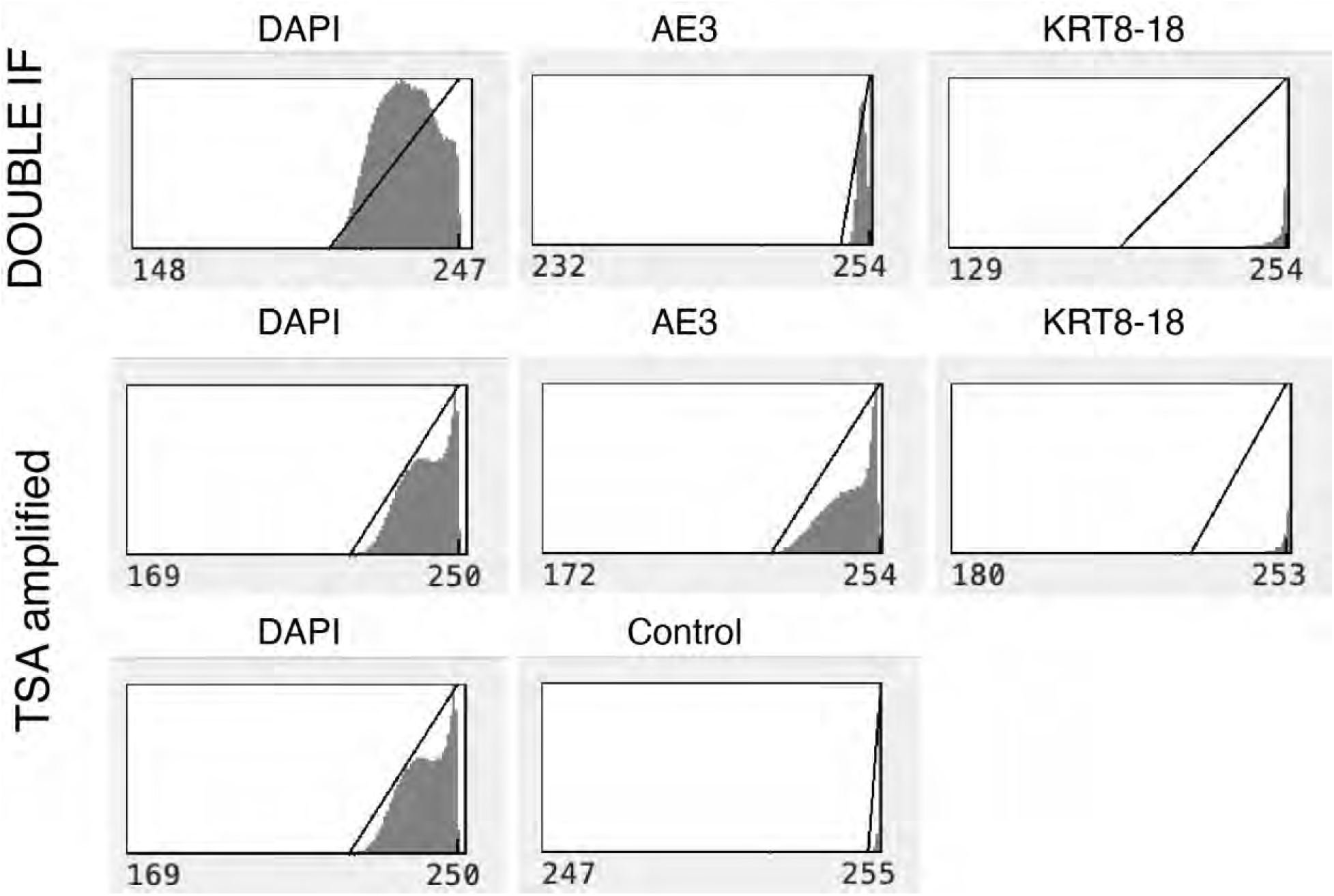
Brightness & Contrast settings for image S8. The Brightness & Contrast settings for the corresponding images of Fig. S8 are shown.

## Notes

https://board.unimib.it/datasets/kmxz7fgydx/1

### Competing Interest Statement

The authors have declared no competing interest.

## References

1. Le, P., N. Ahmed, and G.W. Yeo, Illuminating RNA biology through imaging. Nature Cell Biology, 2022. 24(6): p. 815–824 doi: 10.1038/s41556-022-00933-9 PMID - 35697782.

2. Mund, A., et al., Deep Visual Proteomics defines single-cell identity and heterogeneity. Nature Biotechnology, 2022. 40(8): p. 1231–1240 doi: 10.1038/s41587-022-01302-5. PMID: 35590073.

3. Svensson, V., R. Vento-Tormo, and S.A. Teichmann, Exponential scaling of single-cell RNA-seq in the past decade. Nat Protoc, 2018. 13(4): p. 599–604 doi: 10.1038/nprot.2017.149. PMID: 29494575.

4. De Smet, F., A. Antoranz Martinez, and F.M. Bosisio, Next-Generation Pathology by Multiplexed Immunohistochemistry. Trends in Biochemical Sciences, 2021. 46(1): p. 80–82 doi: 10.1016/j.tibs.2020.09.009. PMID: 33097382.

5. Taube, J.M., et al., The Society for Immunotherapy of Cancer statement on best practices for multiplex immunohistochemistry (IHC) and immunofluorescence (IF) staining and validation. J Immunother Cancer, 2020. 8(1)10.1136/jitc-2019-000155. PMID: 32414858.

6. Herzenberg, L.A., et al., Interpreting flow cytometry data: a guide for the perplexed. Nature Immunology, 2006. 7(7): p. 681–685 doi: 10.1038/ni0706-681. PMID: 16785881.

7. Olsen, L.R., et al., The anatomy of single cell mass cytometry data. Cytometry A, 2019. 95(2): p. 156–172 doi: 10.1002/cyto.a.23621. PMID: 30277658.

8. Mair, F., et al., The end of gating? An introduction to automated analysis of high dimensional cytometry data. European Journal of Immunology, 2016. 46(1): p. 34–43 doi: 10.1002/eji.201545774. PMID: 26548301.

9. van Gijlswijk, R.P., et al., Fluorochrome-labeled tyramides: use in immunocytochemistry and fluorescence in situ hybridization. The Journal of Histochemistry and Cytochemistry, 1997. 45(3): p. 375–382 doi. PMID: 9071319.

10. Aeffner, F., et al., Introduction to Digital Image Analysis in Whole-slide Imaging: A White Paper from the Digital Pathology Association. J Pathol Inform, 2019. 10: p. 9 doi: 10.4103/jpi.jpi_82_18. PMID: 30984469.

11. Clarke, Z.A., et al., Tutorial: guidelines for annotating single-cell transcriptomic maps using automated and manual methods. Nature Protocols, 2021. 16(6): p. 2749–2764 doi: 10.1038/s41596-021-00534-0. PMID: 34031612.

12. Heumos, L., et al., Best practices for single-cell analysis across modalities. Nature Reviews Genetics, 2023. p. 1–23 doi: 10.1038/s41576-023-00586-w.

13. Ljosa, V. and A.E. Carpenter, Introduction to the quantitative analysis of two-dimensional fluorescence microscopy images for cell-based screening. PLoS computational biology, 2009. 5(12): p. e1000603 doi: 10.1371/journal.pcbi.1000603. PMID: 20041172.

14. Jamali, N., et al., 2020 BioImage Analysis Survey: Community experiences and needs for the future. Biol Imaging, 2022. 11010.1017/S2633903X21000039. PMID: 35387317.

15. Czerniak, B., et al., Quantitation of oncogene products by computer-assisted image analysis and flow cytometry. The Journal of Histochemistry and Cytochemistry, 1990. 38(4): p. 463–466 doi. PMID: 1969431.

16. Ma, J. and B. Wang, Towards foundation models of biological image segmentation. Nature Methods, 2023. 20(7): p. 953–955 doi: 10.1038/s41592-023-01885-0.

17. Amitay, Y., et al., CellSighter: a neural network to classify cells in highly multiplexed images. Nature Communications, 2023. 14(1): p. 4302 doi: 10.1038/s41467-023-40066-7.

18. Carpenter, A.E., B.A. Cimini, and K.W. Eliceiri, Smart microscopes of the future. Nature Methods, 2023. 20(7): p. 962–964 doi: 10.1038/s41592-023-01912-0.

19. Dall’Olio, L., et al., BRAQUE: Bayesian Reduction for Amplified Quantization in UMAP Embedding. Entropy (Basel), 2023. 25(2)10.3390/e25020354. PMID: 36832720.

20. Bolognesi, M.M., et al., Multiplex Staining by Sequential Immunostaining and Antibody Removal on Routine Tissue Sections. J Histochem Cytochem, 2017. 65(8): p. 431–444 doi: 10.1369/0022155417719419. PMID: 28692376.

21. Bolognesi, M.M., et al., Antibodies validated for routinely processed tissues stain frozen sections unpredictably. Biotechniques, 2021. 70(3): p. 137–148 doi: 10.2144/btn-2020-0149. PMID: 33541132.

22. Cattoretti, G., et al. Multiple Iterative Labeling by Antibody Neodeposition (MILAN). Protocol Exchange 2019 September 25, 2019; Available from: https://protocolexchange.researchsquare.com/article/nprot-7017/v5.

23. Mascadri, F., et al., Background-free Detection of Mouse Antibodies on Mouse Tissue by Anti-isotype Secondary Antibodies. J Histochem Cytochem, 2021. 69(8): p. 535–541 doi: 10.1369/00221554211033239. PMID: 34282664.

24. Mascadri, F., et al., Rejuvenated Vintage Tissue Sections Highlight Individual Antigen Fate During Processing and Long-term Storage. J Histochem Cytochem, 2021. 69(10): p. 659–667 doi: 10.1369/00221554211047287. PMID: 34541944.

25. Schindelin, J., et al., Fiji: an open-source platform for biological-image analysis. Nature methods, 2012. 9(7): p. 676–682 doi: 10.1038/nmeth.2019. PMID: 22743772.

26. Ruifrok, A.C. and D.A. Johnston, Quantification of histochemical staining by color deconvolution. Analytical and quantitative cytology and histology / the International Academy of Cytology [and] American Society of Cytology, 2001. 23(4): p. 291–299 doi. PMID: 11531144.

27. Bankhead, P., et al., QuPath: Open source software for digital pathology image analysis. Scientific reports, 2017. 7(1): p. 16878 doi: 10.1038/s41598-017-17204-5. PMID: 29203879.

28. Pachitariu, M. and C. Stringer, Cellpose 2.0: how to train your own model. Nature methods, 2022. 19(12): p. 1634–1641 doi: 10.1038/s41592-022-01663-4. PMID: 36344832.

29. Hickey, J.W., et al., Strategies for Accurate Cell Type Identification in CODEX Multiplexed Imaging Data. Front Immunol, 2021. 12: p. 727626 doi: 10.3389/fimmu.2021.727626. PMID: 34484237.

30. Zhang, W., et al., Identification of cell types in multiplexed in situ images by combining protein expression and spatial information using CELESTA. Nature methods, 2022. 19(6): p. 759–769 doi: 10.1038/s41592-022-01498-z. PMID: 35654951.

31. Kimpe, T. and T. Tuytschaever, Increasing the number of gray shades in medical display systems--how much is enough? Journal of digital imaging, 2007. 20(4): p. 422–432 doi: 10.1007/s10278-006-1052-3. PMID: 17195900.

32. Kreit, E., et al., *Biological versus electronic adaptive coloration: how can one inform the other?* Journal of the Royal Society, Interface, 2013. 10(78): p. 20120601 doi: 10.1098/rsif.2012.0601. PMID: 23015522.

33. Aeffner, F., et al., The Gold Standard Paradox in Digital Image Analysis: Manual Versus Automated Scoring as Ground Truth. Arch Pathol Lab Med, 2017. 141(9): p. 1267–1275 doi: 10.5858/arpa.2016-0386-RA. PMID: 28557614.

34. Hötzel, K.J., et al., Synthetic Antigen Gels as Practical Controls for Standardized and Quantitative Immunohistochemistry. Journal of Histochemistry & Cytochemistry, 2019. 67(5): p. 309–334 doi: 10.1369/0022155419832002. PMID: 30879407.

35. Berry, S., et al., Analysis of multispectral imaging with the AstroPath platform informs efficacy of PD-1 blockade. Science, 2021. 372(6547)10.1126/science.aba2609. PMID: 34112666.

36. Kennedy-Darling, J., et al., Highly multiplexed tissue imaging using repeated oligonucleotide exchange reaction. European Journal of Immunology, 2021. 51(5): p. 1262–1277 doi: 10.1002/eji.202048891. PMID: 33548142.

37. Woodcock-Mitchell, J., et al., Immunolocalization of keratin polypeptides in human epidermis using monoclonal antibodies. The Journal of cell biology, 1982. 95(2 Pt 1): p. 580–588 doi. PMID: 6183275.

38. Linden, M.D. and R.J. Zarbo, Cytokeratin immunostaining patterns of benign, reactive lymph nodes: applications for the evaluation of sentinel lymph node specimen. Applied Immunohistochemistry & Molecular Morphology, 2001. 9(4): p. 297–301 doi: 10.1097/00129039-200112000-00002. PMID: 11759054.

39. Franke, W.W. and R. Moll, Cytoskeletal components of lymphoid organs. I. Synthesis of cytokeratins 8 and 18 and desmin in subpopulations of extrafollicular reticulum cells of human lymph nodes, tonsils, and spleen. Differentiation; research in biological diversity, 1987. 36(2): p. 145–163 doi. PMID: 2452110.

40. Argentieri, M.C., et al., Antibodies are forever: a study using 12-26-year-old expired antibodies. Histopathology, 2013. 63(6): p. 869–876 doi: 10.1111/his.12225. PMID: 24102865.

41. Furia, L., et al., Automated multimodal fluorescence microscopy for hyperplex spatial- proteomics: Coupling microfluidic-based immunofluorescence to high resolution, high sensitivity, three-dimensional analysis of histological slides. Front Oncol, 2022. 12: p. 960734 doi: 10.3389/fonc.2022.960734. PMID: 36313693.

42. Saeys, Y., S. Van Gassen, and B.N. Lambrecht, Computational flow cytometry: helping to make sense of high-dimensional immunology data. Nature Reviews Immunology, 2016. 16(7): p. 449–462 doi: 10.1038/nri.2016.56. PMID: 27320317.

43. Segebarth, D., et al., On the objectivity, reliability, and validity of deep learning enabled bioimage analyses. Elife, 2020. 910.7554/eLife.59780. PMID: 33074102.

44. Phillips, D., et al., Highly Multiplexed Phenotyping of Immunoregulatory Proteins in the Tumor Microenvironment by CODEX Tissue Imaging. Frontiers in Immunology, 2021. 12: p. 687673 doi: 10.3389/fimmu.2021.687673 PMID - 34093591.

45. Hickey, J.W., et al., Strategies for Accurate Cell Type Identification in CODEX Multiplexed Imaging Data. Frontiers in immunology, 2021. 12: p. 727626 doi: 10.3389/fimmu.2021.727626 PMID - 34484237.

46. Jarosch, S., et al., Multiplexed imaging and automated signal quantification in formalin- fixed paraffin-embedded tissues by ChipCytometry. Cell Rep Methods, 2021. 1(7): p. 100104 doi: 10.1016/j.crmeth.2021.100104. PMID: 35475000.

47. Stoltzfus, C.R., et al., Multi-Parameter Quantitative Imaging of Tumor Microenvironments Reveals Perivascular Immune Niches Associated With Anti-Tumor Immunity. Frontiers in immunology, 2021. 12: p. 726492 doi: 10.3389/fimmu.2021.726492. PMID: 34421928.

48. Schürch, C.M., et al., Coordinated Cellular Neighborhoods Orchestrate Antitumoral Immunity at the Colorectal Cancer Invasive Front. Cell, 2020. 182(5): p. 1341–1359.e19 doi: 10.1016/j.cell.2020.07.005. PMID: 32763154.

49. Hewitt, S.M., et al., Controls for immunohistochemistry: the Histochemical Society’s standards of practice for validation of immunohistochemical assays. J Histochem Cytochem, 2014. 62(10): p. 693–7 doi: 10.1369/0022155414545224. PMID: 25023613.

50. Chester, C. and H.T. Maecker, Algorithmic Tools for Mining High-Dimensional Cytometry Data. The Journal of Immunology, 2015. 195(3): p. 773–779 doi: 10.4049/jimmunol.1500633. PMID: 26188071.

51. Edfors, F., et al., Enhanced validation of antibodies for research applications. Nat Commun, 2018. 9(1): p. 4130 doi: 10.1038/s41467-018-06642-y. PMID: 30297845.

52. Sulczewski, F.B., et al., Transitional dendritic cells are distinct from conventional DC2 precursors and mediate proinflammatory antiviral responses. Nature Immunology, 2023. 24(8): p. 1265–1280 doi: 10.1038/s41590-023-01545-7.

53. Girard, M., et al., Type I interferons drive the maturation of human DC3s with a distinct costimulatory profile characterized by high GITRL. Science Immunology, 2020. 5(53): p. eabe0347 doi: 10.1126/sciimmunol.abe0347. PMID: 33188059.

54. Yin, X., et al., Human Blood CD1c +Dendritic Cells Encompass CD5 highand CD5 lowSubsets That Differ Significantly in Phenotype, Gene Expression, and Functions. The Journal of Immunology, 2017. 198(4): p. 1553–1564 doi: 10.4049/jimmunol.1600193.

55. Villani, A.-C., et al., *Single-cell RNA-seq reveals new types of human blood dendritic cells, monocytes, and progenitors.* Science (New York, NY), 2017. 356(6335)10.1126/science.aah4573. PMID: 28428369.

56. See, P., et al., Mapping the human DC lineage through the integration of high- dimensional techniques. Science (New York, NY), 2017. 356(6342): p. eaag3009 doi: 10.1126/science.aag3009. PMID: 28473638.

57. Pasqual, G., et al., Monitoring T cell-dendritic cell interactions in vivo by intercellular enzymatic labelling. Nature Publishing Group, 2018. 553(7689): p. 496–500 doi: 10.1038/nature25442. PMID: 29342141.

58. Leylek, R., et al., Integrated Cross-Species Analysis Identifies a Conserved Transitional Dendritic Cell Population. Cell reports, 2019. 29(11): p. 3736–3750.e8 doi: 10.1016/j.celrep.2019.11.042. PMID: 31825848.

59. Wood, G.S. and P.S. Freudenthal, CD5 monoclonal antibodies react with human peripheral blood dendritic cells. The American journal of pathology, 1992. 141(4): p. 789–795 doi. PMID: 1384337.

60. Cattoretti, G., et al., Nuclear and cytoplasmic AID in extrafollicular and germinal center B cells. Blood, 2006. 107(10): p. 3967–3975 doi: 10.1182/blood-2005-10-4170. PMID: 16439679.

61. Willenbrock, K., et al., The expression of activation induced cytidine deaminase in follicular lymphoma is independent of prognosis and stage. Histopathology, 2009. 54(4): p. 509–512 doi: 10.1111/j.1365-2559.2009.03241.x. PMID: 19309412.

62. Ehrhardt, G.R.A., et al., Discriminating gene expression profiles of memory B cell subpopulations. The Journal of experimental medicine, 2008. 205(8): p. 1807–1817 doi: 10.1084/jem.20072682. PMID: 18625746.

63. Kang, Z., et al., Tribus: Semi-automated discovery of cell identities and phenotypes from multiplexed imaging and proteomic data. 2024.10.1101/2024.03.13.584767.

64. Rogalla, S., et al., Automated Spatial Omics Landscape Analysis Approach Reveals Novel Tissue Architectures in Ulcerative Colitis. 2024.10.21203/rs.3.rs-3965505/v1.

65. Wang, K., et al., Comparative analysis of dimension reduction methods for cytometry by time-of-flight data. Nat Commun, 2023. 14(1): p. 1836 doi: 10.1038/s41467-023-37478-w. PMID: 37005472.

66. Amir, E.-a.D., et al., viSNE enables visualization of high dimensional single-cell data and reveals phenotypic heterogeneity of leukemia. Nature Biotechnology, 2013. 31(6): p. 545–552 doi: 10.1038/nbt.2594 PMID - 23685480.

67. Geuenich, M.J., et al., Automated assignment of cell identity from single-cell multiplexed imaging and proteomic data. Cell systems, 2021. 12(12): p. 1173–1186.e5 doi: 10.1016/j.cels.2021.08.012. PMID: 34536381.

## SUPPLEMENTARY REFERENCES

68. Li, C., S. Ziesmer, and O. Lazcano-Villareal, Use of azide and hydrogen peroxide as an inhibitor for endogenous peroxidase in the immunoperoxidase method. The Journal of Histochemistry and Cytochemistry, 1987. 35(12): p. 1457–1460 doi. PMID: 2824601.

69. Manzoni, M., et al., The Adaptive and Innate Immune Cell Landscape of Uterine Leiomyosarcomas. Scientific reports, 2020. 10(1): p. 702–10 doi: 10.1038/s41598-020-57627-1. PMID: 31959856.

70. Liechti, T., et al., An updated guide for the perplexed: cytometry in the high-dimensional era. Nature Immunology, 2021. p. 1–8 doi: 10.1038/s41590-021-01006-z. PMID: 34489590.

